# A cortical hierarchical mechanism for subjective visual experience during working memory

**DOI:** 10.64898/2026.04.08.717338

**Authors:** Tingting Wu, Qing Yu

## Abstract

Working memory (WM) provides a mental sketchpad for simulating sensory experiences associated with mnemonic content. A critical unresolved question is how the brain generates the subjective sensory experience of these internal simulations. Here, we studied individuals with aphantasia, who report a subjective inability to generate visual imagery despite normal WM performance, providing a unique dissociation between objective WM performance and conscious subjective experience. During functional MRI, typical imagers and aphantasic individuals generated and maintained WM content from distinct sources (perception, imagery, or illusion) at a specific spatial location. Time-resolved multivariate analyses revealed a cortical hierarchical mechanism underlying the emergence of subjective visual experience in WM. Rather than dysfunctions in a single brain region, we identified a breakdown in feedback along a cortical pathway from the intraparietal sulcus (IPS) through area V3AB to the early visual cortex (EVC) in aphantasia. The IPS showed impaired imagery generation signals and reduced functional connectivity with EVC, whereas the EVC exhibited a general deficit in feedback independent of voluntary control. These impairments together led to weakened and unstable delay-period representations and precluded the transition from visual to mnemonic codes along the hierarchy. Notably, this network mechanism was specific to imagery at contralateral spatial locations, as ipsilateral IPS maintained compensatory, non-visual representations to support successful task performance without accompanying subjective experience. These results collectively demonstrate that the cortical hierarchy from IPS to EVC is critical for conscious mental visualization of memory content during WM maintenance.

## Introduction

Working memory (WM) provides a mental sketchpad for temporarily maintaining and manipulating information in the absence of external stimulation [1]. This capacity is crucial for mental imagery and simulation, such as visualizing details of a past experience or simulating possible future scenarios, which are key processes for constructing knowledge of the world according to theories of embodied cognition [2]. Converging neuroimaging evidence has revealed that, when holding specific WM content in mind, content-specific WM representations are observed across a distributed cortical network, including frontal, parietal, and sensory regions [3–6]. These representations have been considered core neural signatures of WM maintenance.

However, most WM research to date has focused on its objective dimension, such as the content of information being remembered. It has remained unclear whether these content-specific representations across cortical regions underlie the subjective sensory experience of WM, for instance, mentally visualizing WM content as if real “seeing.” Such conscious visualization may be critical for high-fidelity WM, which requires the maintenance of fine-grained visual details including spatial locations and visual features [7,8]. The sensory recruitment hypothesis of WM proposes that sensory cortices serve as storage sites for WM content to support precise WM of sensory details [9,10]. In support of this view, the visual cortex, in particular the early visual cortex (EVC), demonstrates persistent and dynamic encoding of mnemonic information that reliably tracks WM precision [11,12], biases [13–15], and task demands [7,8,16].

Although EVC may provide a basis for mental simulation of subjective visual experience, direct evidence supporting this view remains limited. This is largely due to a lack of reliable measures of subjective experience compared to objective task performance. Recent advances in aphantasia research offer a unique and compelling opportunity to test this question. Individuals with aphantasia report an inability to voluntarily generate and sustain visual imagery in the absence of external input [17,18]. Although they are aware of the content they intend to visualize and are able to perform cognitive tasks such as WM [19–21], they subjectively experience no conscious “seeing” of WM content during WM delay. The fact that aphantasics can perform WM tasks suggests a dissociation between WM maintenance and the subjective visual experience associated with this process. Comparing neural activity between aphantasics and typical imagers can therefore help clarify the neural underpinnings of subjective visual experience during WM maintenance.

A critical question, then, is whether aphantasia results from dysfunction of specific brain regions like the EVC [22,23] or a broader breakdown in network-level functions. Recent evidence suggests that content-specific representations in EVC alone may not support stable maintenance in a changing environment, as they are vulnerable to distractors [15,24,25]. Moreover, delay-period representations in EVC are still observed during visual imagery or memory retrieval when external stimulation is absent [26–31], and are localized to the feedback layers of EVC by layer-specific functional MRI (fMRI) [32,33]. These results suggest that delay-period EVC representations likely reflect feedback activity from higher-order regions rather than residual sensory-driven responses carried over from encoding.

The frontal and parietal cortices are potential sources of feedback signals to EVC [27,34–38]. The frontal cortex, particularly the PFC, has long been implicated in exerting top-down control over stimulus representations in sensory regions, including the EVC [9,39]. The parietal cortex, by contrast, may play a more specific role in generating imagery or WM content [28,40,41]. Moreover, the functional coupling between frontoparietal and visual cortices has been shown to underlie successful WM performance [42,43] and imagery vividness [37]. Despite the established link, the precise nature of these feedback signals and the specific mechanisms by which they support WM remain to be further elucidated.

Based on the above, if impaired content-specific representation during delay is observed only in the EVC of aphantasia, it would suggest that diminished subjective visual experience is due to local dysfunction in the EVC. Alternatively, if both EVC representations and the putative source signals in the frontoparietal cortex are disrupted, addressing the information flow between these regions would be crucial for determining whether aphantasia is due to a broader breakdown across the cortical hierarchy from higher-order frontoparietal regions to EVC.

To investigate the neural mechanisms underlying subjective visual experience during WM in a time-resolved manner, we designed three WM tasks, each involving a delayed-recall trial structure but differing in how WM stimulus was delivered [28]. The first was a perception-based WM task, in which participants viewed a visual stimulus to be remembered. This task ensured comparable feedforward input to EVC between aphantasics and typical imagers. The second was an imagery-based WM task that began with a symbolic cue instructing which stimulus to imagine during WM. Critically, because direct retinal input could not account for conscious visual experience in this task, feedback signals from higher-order cortex were required even during the encoding stage. This design rendered the subjective visual experience of typical imagers and aphantasics distinct across tasks and stages: during the sample period, aphantasics subjectively experienced a failure in voluntary imagery generation in the imagery task but reported normal visual perception in the perception task. During the delay period, however, they experienced an inability to sustain mental imagery in both tasks. This dissociation allowed us to separately examine the neural correlates of imagery generation and maintenance. In addition, we included a third illusion-based WM task, in which participants viewed an illusory stimulus producing distinct conscious visual experiences despite near-identical visual input. Notably, illusion has been conceptualized as “involuntary imagery,” differing from voluntary imagery in the degree of intentionality [35,44]. This offers an opportunity to test whether deficits in aphantasia are specific to voluntary imagery control. Together, this design allowed us to investigate whether feedforward and feedback activity differ between typical imagers and aphantasics across brain regions, when these differences, if exist, emerge across different task stages, and whether they can account for group differences in subjective visual experience.

## Results

### Aphantasics successfully performed working memory tasks but with worse performance than typical imagers

We conducted a series of questionnaires and interviews to ensure that the recruited aphantasic participants truly lacked visual imagery. All aphantasic participants self-reported low scores on the Vividness of Visual Imagery Questionnaire (VVIQ) (16.2 ± 0.7 in aphantasics vs. 61.8 ± 9.1 in typical imagers, Figure S1A). Self-reports on other imagination-related questionnaires also indicated an absence of visualization during mental imagery in these participants (Figure S1B). In addition to impaired visual imagery, most participants also reported less spontaneous use of visual imagery (Figure S1C), and experienced varying degrees of auditory, olfactory, gustatory, and tactile imagery deficits (Figure S1D-F and Table S1). Moreover, while all participants reported having dreams, only a few (5 out of 17) considered visual images during their dreams to be entirely normal. These results are in line with previous reports on the heterogeneity of aphantasia [45,46]. A complete summary of participants’ self-reports can be found in Table S1.

Inside an MRI scanner, both typical imagers (n = 17) and aphantasics (n = 17) completed three WM tasks that differed only during the sample period (Figure 1A) [28]. In the perception-based WM task (perception), participants viewed a Gabor pattern moving back and forth along a tilted path (left or right) in the right visual periphery. In the imagery-based WM task (imagery), a symbolic cue appeared at fixation, instructing participants to imagine a Gabor moving in the cued direction at the same peripheral location as the perception task. In the illusion-based WM task (illusion), participants viewed a Gabor moving vertically, with its internal texture drifting leftward or rightward, inducing a strong illusory percept of left- or right-tilted motion [47,48]. During a prolonged delay period, participants were instructed to continue maintaining the moving Gabor along the same tilted path at the same location. Subsequently, they reported the memorized motion path as accurately as possible on an orientation wheel. Participants were instructed to maintain fixation throughout the tasks. Note that motion paths were consistent across tasks. Because illusion size (i.e., the degree of tilt) varied across participants, individuals’ illusion sizes were measured prior to scanning, and the degree of tilt on perception trials was tailored accordingly (see methods). Likewise, on imagery trials, participants were asked to imagine a moving Gabor with motion paths matched to the perception and illusion trials. Participants were unaware of the illusion (i.e., they believed that all illusion trials were perception trials), as confirmed by a post-experiment interview.

**Figure 1.**
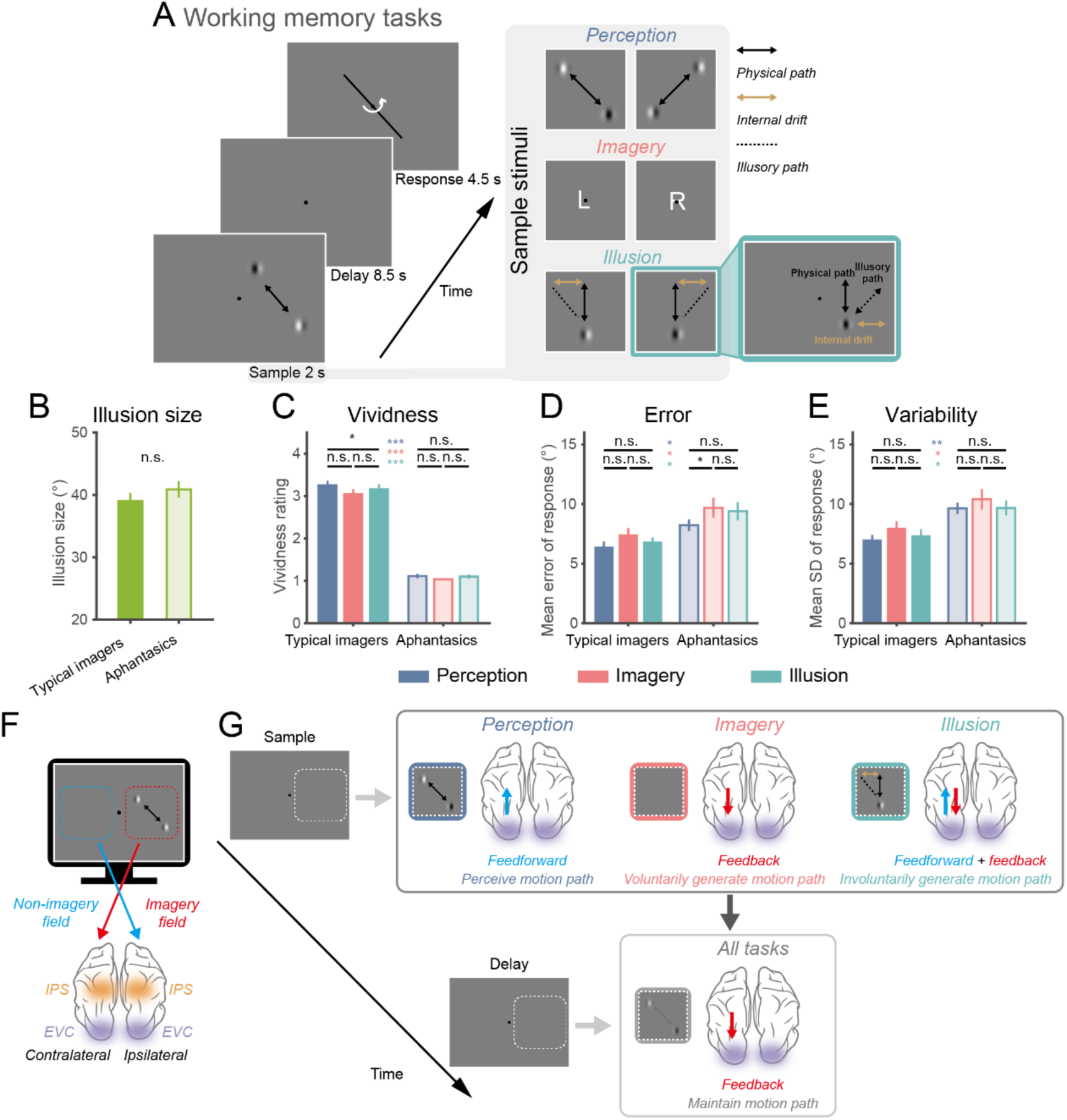
Experimental paradigm and behavioral performance. (A) WM tasks in the MRI scanner. Participants performed three WM tasks, differing only in the stimulus presented during the sample period. In the perception task, a Gabor patch moved back and forth along a left- or right-tilted path. In the imagery task, a symbolic cue indicated whether a Gabor moving along left- or right-tilted path should be imagined. In the illusion task, a Gabor with an internal texture drifting leftward or rightward moved back and forth vertically, creating the illusion of left- or right-tilted motion. During the subsequent delay, participants mentally maintained or imagined a dynamic scene in which the Gabor moved back and forth along the same direction, in the same position as the originally presented Gabor. After the delay, participants reported the maintained motion direction. (B) Measured illusion size. Illusion size measured prior to the main task (inside the MRI scanner) for typical imagers (n = 17) and aphantasics (n = 17), which was used to parameterize subsequent WM tasks. Colored bars indicate group average, error bars represent ±1 SEM. (C) Vividness ratings for delay-period experience. Ratings for each WM task were collected in a separate behavioral training session from typical imagers (n = 16) and aphantasics (n = 17). Black asterisks indicate significant differences across tasks within the same group; colored asterisks indicate significant differences between groups for the same task. Colored bars represent group means, with error bars representing ±1 SEM. (D) Mean recall error for WM reports. Conventions are the same as in (C). (E) Response variability for WM reports, calculated as the standard deviation (SD) of responses. Conventions are the same as in (C). Significance levels: ***, *p* < 0.001; **, *p* < 0.01; *, *p* < 0.05; n.s., *p* ≥ 0.05. All p-values were FDR-corrected. (F) Definition of imagery and non-imagery fields. The side of the screen where sample stimuli were presented was defined as the imagery field, whereas the opposite side was defined as the non-imagery field. For neural analyses, brain regions in the hemisphere contralateral to the imagery field were considered to be directly related to subjective visual experience (where present), whereas ipsilateral regions were not. (G) Schematic illustration of hypothetical feedforward and feedback activity during sample and delay periods across three tasks. During the sample period, the perception task primarily involved feedforward activity in the EVC driven by the moving Gabor. The imagery task, by contrast, required voluntary generation of the motion path in higher-order cortices without direct stimulation of the EVC. The illusion task involved both feedforward activity driven by the physical Gabor and feedback activity from higher-order cortices for the involuntary generation of the illusory motion path. During the delay period, no external visual stimulation was provided, and EVC was predominantly driven by feedback activity across all tasks.

First, we analyzed vividness ratings of WM content during the delay period (Figure 1C), collected during a separate behavioral training session. Using a 4-point scale (1 = no imagery, 4 = vivid “as if seen” imagery), aphantasics (perception = 1.1±0.2, imagery = 1.0±0.1, illusion = 1.1±0.2) reported significantly lower vividness across all three tasks compared to typical imagers (perception = 3.3±0.4, imagery = 3.1±0.5, illusion = 3.2±0.5; *t*s > 15.49, *p*s < 0.001; all p-values were FDR corrected in this and all subsequent analyses unless specified). There were no significant differences between tasks within each group (*t*s < 2.0, *p*s > 0.113), except that perception was rated as slightly more vivid than illusion in typical imagers (*t*(15) = 3.1, *p* = 0.022). This confirmed that aphantasics’ subjective conscious experience during the delay period was markedly different from that of typical imagers, consistent with their reported absence of visual imagery. Note that vividness ratings from typical imagers were obtained from a separate group of participants (n = 16) who did not participate in the fMRI scans.

Despite this marked difference in subjective experience, both groups reported a large illusion size around 40° (typical imagers: 39.0±5.3°, aphantasics: 40.9±5.6°, *t*(32) = 1.0, *p* = 0.335), demonstrating comparable illusory perception (Figure 1B). This suggests that, aphantasics’ subjective visual experience during the generation of involuntary imagery, at least for motion illusions, remains intact, forming the basis for further comparisons based on illusory motion paths. In terms of behavioral performance during WM, while aphantasics successfully completed the WM tasks with a mean recall error of 9.1±3.0° (perception = 8.2±2.0°, imagery = 9.7±3.6°, illusion = 9.4±3.3°), their response variability (perception = 9.6±2.0°, imagery = 10.4±3.5°, illusion = 9.7±2.6°) and recall error were significantly larger than typical imagers (recall error: perception = 6.4±2.0°, imagery = 7.4±2.4°, illusion = 6.8±1.7°, *t*s > 2.0, *p*s < 0.047; response variability: perception = 6.9±1.8°, imagery = 7.9±2.4°, illusion = 7.3±2.2°, *t*s > 2.1, *p*s < 0.039; Figure 1D-E). This indicates that although aphantasics can perform WM tasks without visual imagery, their performance is less accurate, likely due to the absence of mental images supporting precise WM of motion paths.

### Absence of imagery dominance in IPS of aphantasics during imagery generation

To investigate how typical imagers and aphantasics encoded WM content at the neural level, we conducted a time-resolved decoding analysis on the direction of motion paths. Because participants perceived and maintained information in the right visual field only, contralateral regions of interest (ROIs) corresponded to the visual field that directly engaged subjective visual experience in typical imagers (i.e., the imagery field), whereas ipsilateral ROIs received no direct external input and did not engage subjective visual experience (i.e., the non-imagery field, Figure 1F). We first focused our analyses on the EVC and the intraparietal sulcus (IPS), which has long been implicated in both WM and mental imagery for motion directions and line orientations [24,25,28,49]. Support vector machine (SVM) decoders were used to discriminate neural activity patterns for leftward versus rightward motion paths, at each time point for each task. This time-resolved approach allowed us to track the temporal unfolding of mental representations for motion paths in specific ROIs. In particular, we could separately examine feedforward versus feedback activity, as well as content generation versus maintenance, across tasks and stages (sample vs. delay periods; Figure 1G).

First, we asked how specific imagery content was generated during the sample period. Unlike the perception task which required minimal voluntary imagery control for perceiving external stimuli during the sample period, successfully generating the cued imagery content did require voluntary control. Comparisons between imagery and perception tasks would therefore identify where in the brain this imagery generation signal originated. We found that, motion paths could be decoded in contralateral EVC and IPS in both tasks for both typical imagers and aphantasics (Figure 2A and 2C). However, the decoding strength differed between tasks. Specifically, previous work from our lab has identified a neural signature in the IPS, the imagery dominance signal, which potentially underlies imagery generation signals of specific content in typical imagers [28]. Imagery dominance was defined as superior neural decoding strength during imagery compared to perception. Likewise, a reverse pattern with superior decoding strength for perception was considered perception dominance. Perception dominance was observed in contralateral EVC (Figure 2B left panel, *p* < 0.001), suggesting dominant sensory-driven signals in this region during the sample period. By contrast, imagery dominance was found in contralateral IPS (Figure 2B left panel, *p* = 0.012), suggesting dominant imagery signals that contributed to imagery generation during the sample period. Extending this analysis to retinotopic ROIs along a posterior-to-anterior cortical gradient (V1 to V3, V3AB, IPS0 to IPS5) revealed a gradual shift from perception dominance in contralateral EVC to imagery dominance in contralateral IPS, with V3AB demonstrating an equilibrium between imagery and perception (Figure 2B right panel). Moreover, this transition from perception dominance in the EVC to imagery dominance in the IPS occurred only in the contralateral but not in the ipsilateral pathway, indicating the location specificity of the effect (Figure S2A and S2B). These results together suggest that imagery dominance develops gradually along the cortical visual hierarchy from EVC to IPS.

**Figure 2.**
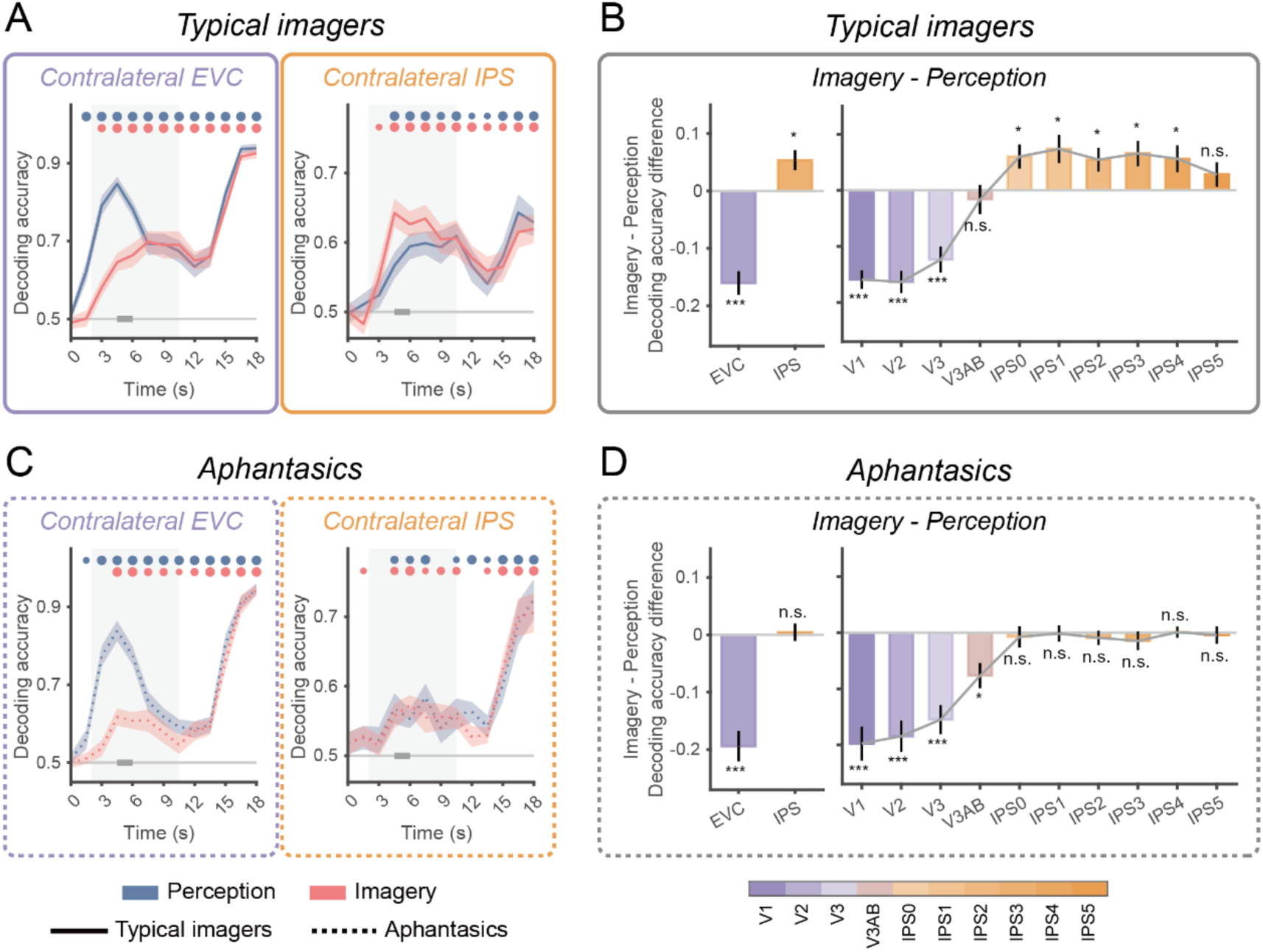
Imagery dominance in contralateral EVC and IPS in typical imagers and aphantasics. (A) Time course of decoding accuracy for perception (blue) and imagery (red) tasks in contralateral EVC (left panel) and IPS (right panel) of typical imagers. Circles on the top denote significant time points. Circle size reflects significance level: large, *p* < 0.001; medium, *p* < 0.01; small, *p* < 0.05. (B) Imagery dominance in typical imagers. Left panel: Imagery dominance in contralateral EVC and IPS during the sample period (time windows indicated by thick gray bars in A), defined as the difference in decoding accuracy between imagery and perception tasks. Error bars indicate ±1 SEM. Asterisks denote significant differences between perception and imagery within the corresponding ROI. Right panel: Imagery dominance across contralateral visual hierarchy (V1 to IPS5), using the same conventions. Significance levels: ***, *p* < 0.001; **, *p* < 0.01; *, *p* < 0.05; n.s., *p* ≥ 0.05. All p-values were FDR-corrected. (C) Same layout and conventions as (A), with results from aphantasics (dashed lines). (D) Imagery dominance in aphantasics. Same layout and conventions as (B).

When applying the same analyses to aphantasics, we found that this imagery dominance disappeared. While contralateral EVC in aphantasics still showed perception-dominant signals (*p* < 0.001; Figure 2C left panel), imagery dominance in contralateral IPS was absent (*p* = 0.778; Figure 2C right panel), as was the posterior-to-anterior gradient (Figure 2D). Ipsilateral ROIs demonstrated a similar null effect overall, although weak imagery dominance was found in the ipsilateral IPS (Figure S2C and S2D), possibly reflecting a compensatory role of ipsilateral IPS in imagery generation in aphantasia. We further confirmed that the absence of imagery dominance was not specific to the right visual field, as similar results were observed when WM was performed in the left visual field in a control experiment (n = 6; Figure S3).

To what extent is this imagery dominance specific to IPS? To address this question, we examined four additional ROIs, the motion-selective MT+, the occipital fusiform gyrus and the temporal occipital fusiform cortex in the ventral visual pathway [50–52], as well as the superior precentral sulcus (sPCS) in the lateral frontal cortex, which has also frequently been implicated in WM [4,6]. We found that none of these ROIs demonstrated a clear imagery dominance effect comparable to the IPS. Imagery dominance was observed in the MT+ of both groups (Figure S4A-B). However, this effect was present not only in the contralateral but also in the ipsilateral MT+, suggesting that imagery dominance in the MT+ may not be directly relevant to subjective visual experience. Motion paths could be decoded from both groups in the occipital fusiform gyrus, but no imagery dominance was found in either hemisphere of either group (Figure S4C-D). The temporal occipital fusiform cortex showed stronger imagery dominance in aphantasia, with no hemispheric difference (Figure S4E-F). In the sPCS, decoding accuracy for perception and imagery remained overall lower, and again no imagery dominance was detected in either hemisphere of either group (Figure S4G-H). These findings further support the conclusion that imagery dominance signal in the contralateral IPS is specifically linked to voluntary imagery generation and its accompanying subjective experience, and that voluntary imagery generation in aphantasics is impaired.

### Impaired feedback to early visual cortex in aphantasia

Having identified impaired imagery generation signals in the IPS during the imagery task, we next examined delay-period representations that reflected WM maintenance activity. Because aphantasics reported a lack of subjective visual experience during delay across all tasks, we compared the decoding strength for motion paths between typical imagers and aphantasia for each task separately, instead of focusing on the differences between tasks within the same group. In contralateral EVC, motion paths could be decoded in all tasks for both typical imagers and aphantasics (Figure 3A). Additionally, no significant difference was found between groups in contralateral EVC during the sample period (4.5 – 6 s) of the perception task (*p* = 0.793), suggesting that aphantasics responded to external visual input similarly to typical imagers, forming normal feedforward representations of motion paths. However, a divergence between groups emerged gradually during the delay period (9 – 10.5 s): aphantasics showed significantly lower decoding accuracy for motion paths (*p* = 0.027), indicating that they were unable to maintain content-specific representations at the same strength as typical imagers. Because delay-period representations rely on feedback signals, this group difference was likely due to impaired feedback to EVC in aphantasia. Similarly weakened representations were also observed in imagery (*p* = 0.013) and illusion (*p* = 0.024) tasks during the delay. Importantly, the divergence between groups in these two tasks emerged earlier than in the perception task (7.5 s for imagery and 6 s for illusion versus 9 s for perception), suggesting that feedback signals were already impaired prior to WM maintenance in aphantasia. Since the generation of illusory motion during the sample period is thought to recruit higher-order cortices [47], these results indicate that the feedback impairment in aphantasia is not limited to WM maintenance but may reflect a general deficit. Importantly, this deficit occurred regardless of whether the imagery process was voluntary (imagery versus illusion), suggesting that it is not dependent on voluntary imagery control.

**Figure 3.**
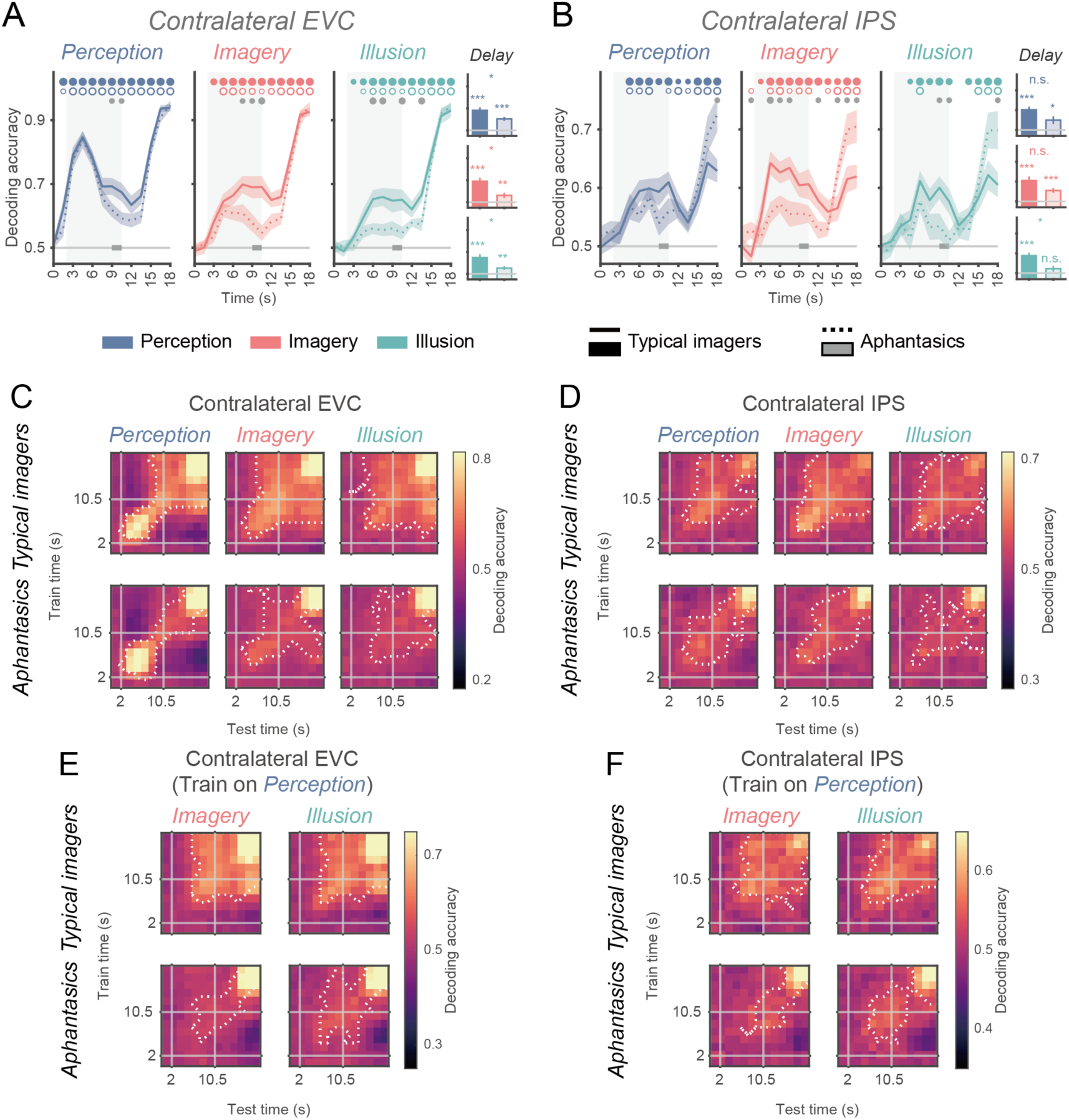
Comparison of motion path decoding during WM tasks between typical imagers and aphantasics in contralateral EVC and IPS. (A) Time course of decoding accuracy in contralateral EVC for perception (blue, left), imagery (red, middle), and illusion (green, right) tasks in typical imagers (solid lines) and aphantasics (dashed lines). The solid gray horizontal line denotes chance-level performance. The gray shaded region indicates the delay period. Thick gray horizontal bars on the chance line mark the time windows used to compute the mean decoding accuracy shown in the bar plots on the right. Circles on the top denote significant time points: filled circles indicate typical imagers, hollow circles indicate aphantasics, and gray circles indicate between-group differences. Circle size reflects significance level: large, *p* < 0.001; medium, *p* < 0.01; small, *p* < 0.05. The three bar plots on the right summarize mean decoding accuracy during the delay period for perception, imagery, and illusion tasks. Solid bars represent typical imagers, hollow bars represent aphantasics. Error bars indicate ±1 SEM. Asterisks above individual bars denote significant decoding accuracy within the corresponding group and task, asterisks between bars indicate significant differences between groups for a given task. Significance levels: ***, *p* < 0.001; **, *p* < 0.01; *, *p* < 0.05; n.s., *p* ≥ 0.05. All p-values were FDR-corrected except those for the time course of between-group differences. (B) Same layout and conventions as (A), with results from contralateral IPS. (C) Cross-temporal generalization matrices in contralateral EVC for perception (left), imagery (middle), and illusion (right) tasks (top row: typical imagers; bottom row: aphantasics). The x- and y-axes denote testing time and training time, respectively. White dashed contours outline clusters exhibiting significant motion path representations after cluster-based multiple-comparison correction. (D) Same layout and conventions as (C), with results from contralateral IPS. (E) Cross-task generalization results for typical imagers (top row) and aphantasics (bottom row) in contralateral EVC. For each group, the classifiers trained on the perception task and tested separately on the imagery task (left) and the illusion task (right). Axes indicate testing time (x-axis) and training time (y-axis). (F) Same layout and conventions as (E), with results from contralateral IPS.

Other key features of delay-period WM representations are their temporal stability [28,53,54] and resemblance to perception [3,25,28,55]. We therefore separately examined whether these features were preserved in aphantasia. First, for each task, we used cross-temporal generalization decoding to assess the stability of time-varying population dynamics for motion path representations. Overall, aphantasics exhibited weaker cross-temporal generalization within each task in contralateral EVC (Figure 3C). Second, we assessed cross-task generalization of neural representations between imagery or illusion trials and perception trials, by training a classifier on perception data and testing it on the other tasks. Again, aphantasics showed reduced shared representations between perception and the two other tasks in contralateral EVC, compared to typical imagers (Figure 3E). These results suggest that, in addition to weaker representations, aphantasics exhibit more temporally variable neural representations across the delay period, which may again reflect impaired feedback in sustaining a coherent and stable visual experience during WM maintenance.

### Content-specific representations are impaired in contralateral IPS but remain intact in ipsilateral IPS

Having demonstrated impaired representations of motion paths in contralateral EVC of aphantasics, we next asked whether similar disruptions exist beyond EVC. We had previously shown impaired imagery generation signals in the IPS. Here, we further investigated whether delay-period IPS representations were also impaired. We found that contralateral IPS also showed reliable decoding of motion paths in both groups (Figure 3B), but the temporal decoding profiles differed from contralateral EVC. In perception and illusion tasks, decoding was weaker in aphantasics; however, the divergence between groups emerged later (for illusion, 9-10.5 s, *p* = 0.014) or was only marginally significant (for perception, 9-10.5 s, *p* = 0.058). This result provides further evidence for impaired feedback to EVC even when higher-order cortices, such as the IPS, retain sufficient information about motion paths during early delay. By contrast, in the imagery task, the divergence emerged during the sample period, even earlier than EVC (4.5 – 6 s, *p* = 0.005), although this difference became slightly weaker during the delay and remained marginally significant (9-10.5 s, *p* = 0.058). This result aligned with the role of IPS in voluntary imagery generation. Additionally, decoding strength in contralateral IPS during the delay period correlated with behavioral response error in both imagery and illusion tasks (Figure S5), suggesting a potential role of IPS delay-period representation in supporting WM maintenance. In summary, while impaired feedback to EVC is general across WM tasks, the demand for voluntary imagery generation engages additional impairment in the IPS. During the delay when external stimulation is absent, this deficit may further affect their ability to sustain visual imagery during WM maintenanc, ultimately leading to diminished conscious visual experience in this process.

Despite marked differences in neural decoding in contralateral EVC and IPS, aphantasics were still able to perform WM tasks. How did they manage this? To address this question, we extended our analyses to ipsilateral ROIs corresponding to the non-imagery field. In ipsilateral IPS (Figure 4B), decoding performance did not significantly differ between groups across tasks (except at 9 s with *p* = 0.046 for the perception task). Moreover, group differences in cross-temporal and cross-task generalization were also less pronounced in ipsilateral IPS (Figure 4D and 4F) compared with contralateral IPS (Figure 3D and 3F). This suggests that, beyond contralateral EVC and IPS, other brain regions such as ipsilateral IPS still represented motion paths normally, possibly in a format not tied to conscious visual imagery. Moreover, this result further supports intact feedforward transmission from contralateral EVC to higher-order cortices, because ipsilateral IPS would otherwise have no source for these signals. In addition, we confirmed that this signal did not originate from ipsilateral EVC either, as decoding performance for motion paths in this region was significantly lower in aphantasics compared to typical imagers regardless of task (Figure 4A; also see Figure 4C and 4E for cross-temporal and cross-task generalization results). Together, these results indicate that aphantasics likely relied on non-visual neural codes, such as those in ipsilateral IPS, to support WM without subjective visual experience.

**Figure 4.**
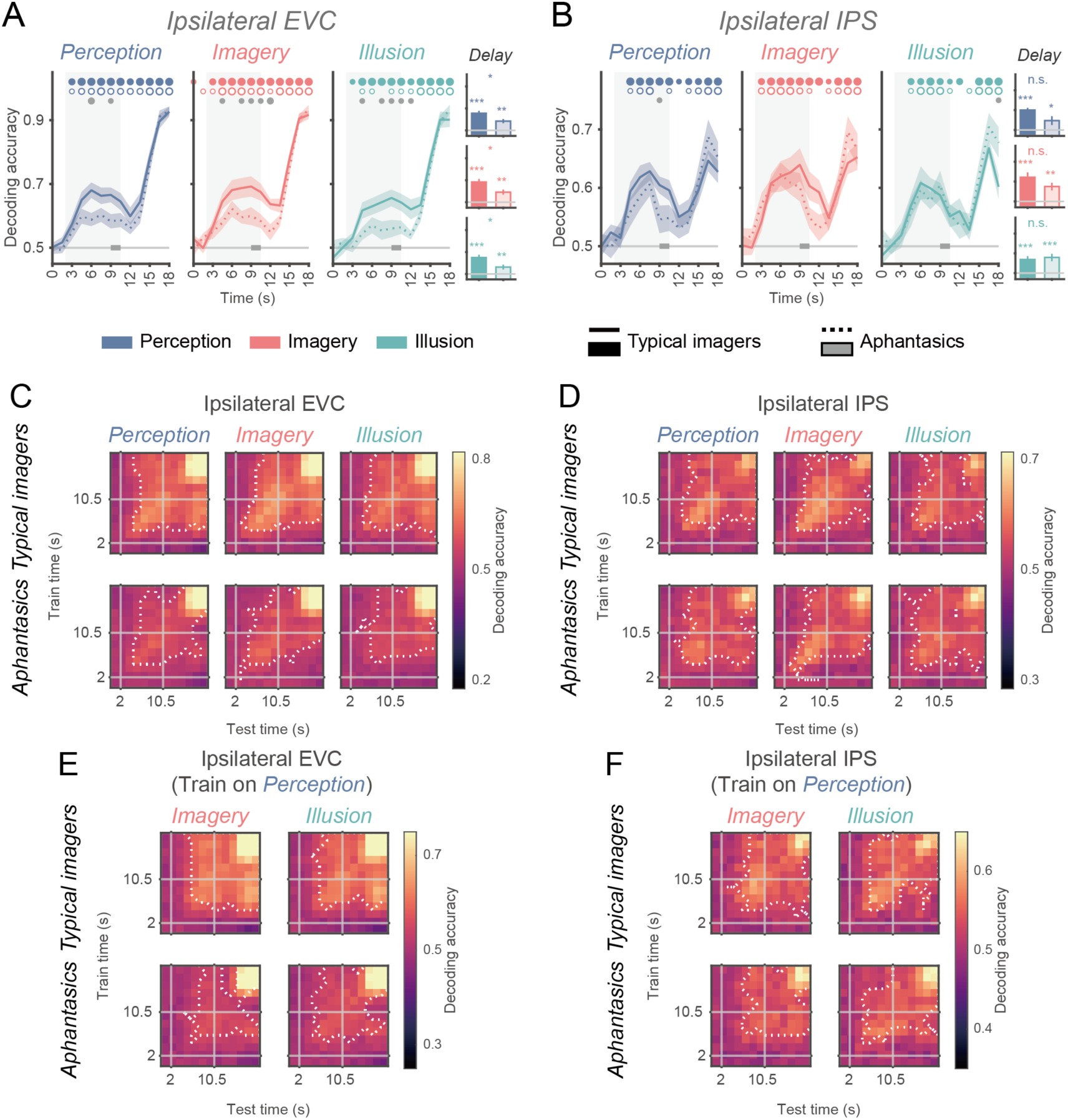
Comparison of motion path decoding during WM tasks between typical imagers and aphantasics in ipsilateral EVC and IPS. (A) Time course of decoding accuracy in ipsilateral EVC for typical imagers (solid lines) and aphantasics (dashed lines), presented separately for perception (blue, left), imagery (red, middle), and illusion (green, right) tasks. The solid gray horizontal line denotes chance-level performance. The gray shaded region indicates the delay period. Thick gray horizontal bars on the chance line mark the time windows used to compute the mean decoding accuracy shown in the bar plots on the right. Circles on the top denote significant time points: filled circles indicate typical imagers, hollow circles indicate aphantasics, and gray circles indicate between-group differences. Circle size reflects significance level: large, *p* < 0.001; medium, *p* < 0.01; small, *p* < 0.05. The three bar plots on the right show mean decoding accuracy during the delay period for perception, imagery, and illusion tasks. Solid bars represent typical imagers, hollow bars represent aphantasics. Error bars indicate ±1 SEM. Asterisks above individual bars denote significant decoding accuracy within the corresponding group and task, asterisks between bars indicate significant differences between groups for a given task. Significance levels: ***, *p* < 0.001; **, *p* < 0.01; *, *p* < 0.05; n.s., *p* ≥ 0.05. All p-values were FDR-corrected except those for time course of between-group differences. (B) Same layout and conventions as (A), with results from ipsilateral IPS. (C) Cross-temporal generalization matrices in ipsilateral EVC for perception (left), imagery (middle), and illusion (right) tasks (top row: typical imagers; bottom row: aphantasic participants). The x- and y-axes represent testing time and training time, respectively. White dashed contours outline clusters exhibiting significant motion path representations after cluster-based multiple-comparison correction. (D) Same layout and conventions as (C), with results from ipsilateral IPS. (E) Cross-task generalization results in ipsilateral EVC for typical imagers (top row) and aphantasics (bottom row). For each group, the classifiers trained on the perception task and tested on the imagery task (left) and the illusion task (right), respectively. Axes indicate testing time (x-axis) and training time (y-axis). (F) Same layout and conventions as (E), with results from ipsilateral IPS.

### A bottleneck for the transition between neural codes around V3AB

Although neural representations for motion paths were degraded in aphantasia, they were nevertheless significant, raising questions of what neural formats these representations might take. To further unpack the nature of the motion path representations, we used representational similarity analysis (RSA) [56] to test two hypothesized models, i.e., whether the representations were better explained by visual or mnemonic processing. The visual representational dissimilarity matrix (RDM) primarily reflected differences in the visual stimuli presented during the sample period (Figure 5A), whereas the mnemonic RDM captured mnemonic differences (left versus right) during the delay period independent of WM tasks (Figure 5B). Partial Spearman correlations were computed to control for collinearity between visual and mnemonic RDMs.

**Figure 5.**
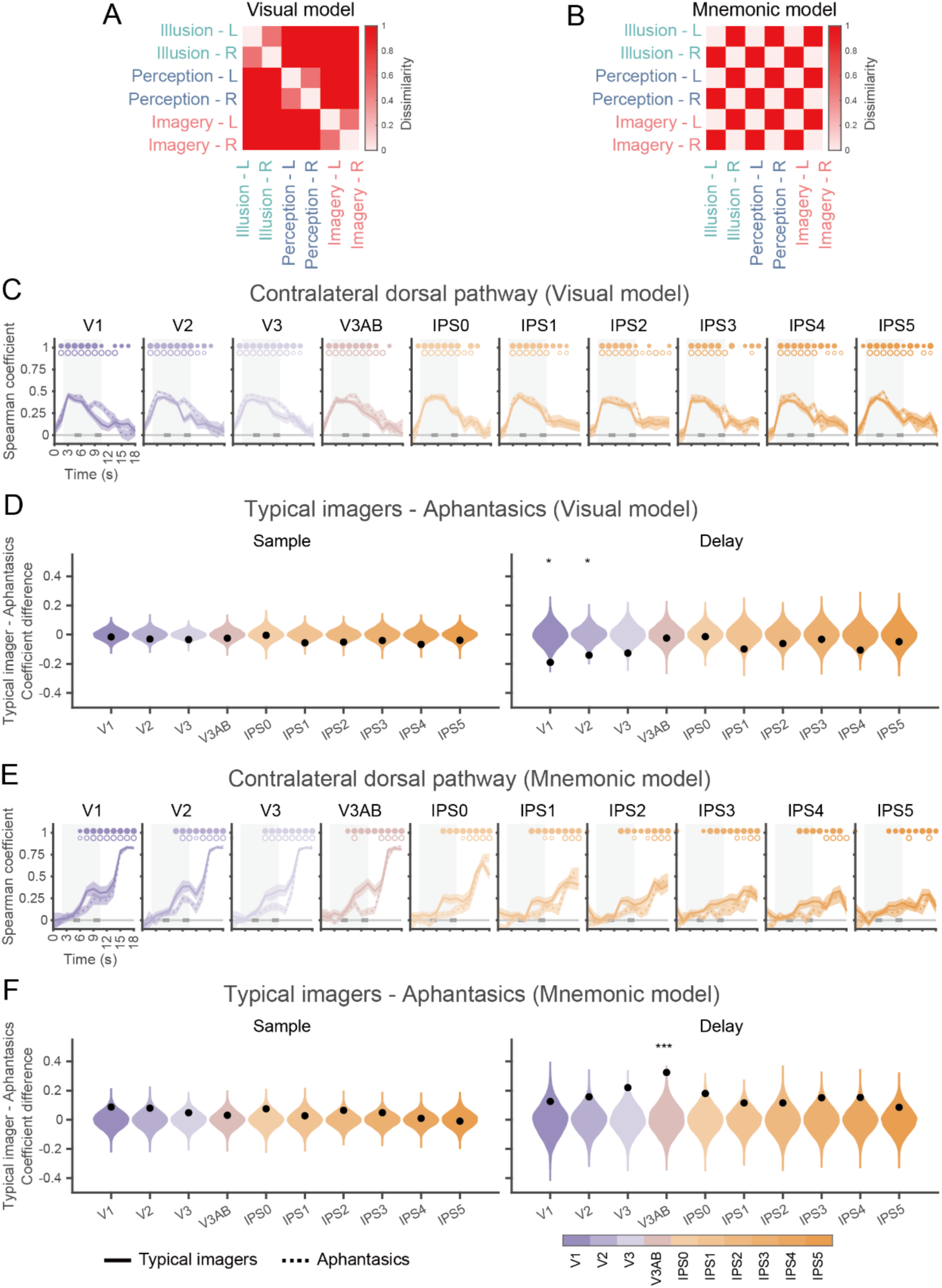
Temporal dynamics of motion-path representational formats from contralateral V1 to IPS5 in typical imagers and aphantasics. (A) Visual model. Hypothesized dissimilarity structure among six motion paths (3 tasks × 2 motion paths) that reflects visual differences during the sample period. Dissimilarity is maximal between motion paths from different tasks, intermediate between different paths within the same task, and minimal between identical paths. (B) Mnemonic model. Hypothesized dissimilarity structure that reflects WM differences during the delay period, defined solely by motion-path identity: dissimilarity is 0 for identical paths and 1 for non-identical paths. (C) Neural similarity to the visual model over time. Spearman correlations between neural RDMs and the visual model are shown as a function of time for contralateral V1 through IPS5 (left to right) in both groups. Solid lines indicate typical imagers, dashed lines indicate aphantasics. Significant time points are marked above each plot (filled circles for typical imagers, hollow circles for aphantasics), with circle size denoting the significance level (large for *p* < 0.001, medium for *p* < 0.01, and small for *p* < 0.05). All p-values were FDR-corrected. (D) Difference in RDM similarity between typical imagers and aphantasics. Left panel: The violin plots depict the null distribution of group differences in Spearman correlation coefficients for the visual model during the sample period (left thick gray bar in C), derived from permutation testing. The black dots denote true group differences. Right panel: Corresponding results for the delay period (right thick gray bar in C). Asterisks statistical significance: *** *p* < 0.001; ** *p* < 0.01; * *p* < 0.05. All p-values were FDR-corrected. (E) Neural similarity to the mnemonic model over time. Same layout and conventions as (C). (F) Same layout and conventions as (D), but with results for the mnemonic model.

In contralateral ROIs, similarity with the visual RDM peaked during the sample period and then declined toward the delay (Figure 5C). In contrast, similarity with the mnemonic RDM gradually increased over time (Figure 5E). Notably, this decline in visual similarity was slower in aphantasics than in typical imagers, particularly in contralateral V1 and V2 during the delay period (Figure 5D). Meanwhile, aphantasics exhibited a slower or even no increase in mnemonic similarity, with the most prominent effect in V3AB during delay (Figure 5F). When using an alternative visual RDM (based on parametric modulation of physical motion-path direction), the overall pattern of results in contralateral ROIs remained largely consistent (Figure S6). In ipsilateral ROIs, both groups showed a gradual decrease in visual similarity and increase in mnemonic similarity over time (Figure S7A and S7C). However, while aphantasics showed a slower decrease in visual similarity from V1 through V3 (Figure S7B), the slower increase in mnemonic similarity was only found in ipsilateral V3 during the sample but not delay period (Figure S7D). These results suggest that, during the delay period, task demands and feedback from IPS likely drove the transition from visual to mnemonic codes in typical imagers, whereas aphantasics continued to represent motion directions differently across tasks when the required feedback was not delivered properly. In addition, V3AB may serve as a bottleneck along the posterior-to-anterior visual hierarchy, where feedback from higher-order cortices, such as the IPS, is blocked in aphantasia.

### Reduced functional connectivity between IPS and visual areas in aphantasia during working memory maintenance

The finding that impaired neural representations of motion paths develop gradually along the posterior-to-anterior dorsal visual hierarchy suggests that the EVC and IPS may be functionally disconnected in aphantasia. To test this directly, we examined functional connectivity between IPS and EVC within the same hemisphere using generalized psychophysiological interaction (gPPI) [57].

Both groups exhibited significant positive functional connectivity between the contralateral EVC and IPS during both the sample (*p*s < 0.047; Figure S8A left panel) and probe (*p*s < 0.007; Figure S8A right panel) periods, with no significant differences between groups (*p*s > 0.143). During the delay period, however, typical imagers showed significant positive connectivity (*p*s < 0.003), whereas aphantasics displayed minimal connectivity (*p*s > 0.061), with a group difference particularly during imagery and illusion tasks (perception: *p* = 0.115; imagery and illusion: *p*s < 0.05; Figure S8A middle panel). A similar pattern was found in the functional connectivity between contralateral IPS and V3AB during the delay (Figure S8C). In contrast, no such difference was observed in ipsilateral connectivity between IPS and EVC (*p*s > 0.481) or between IPS and V3AB (*p*s > 0.118; Figure S8B and S8D). These results suggest a lack of direct functional cross-talk between contralateral regions specifically during the delay in aphantasia. Combined, these findings offer converging evidence that the IPS likely provides the feedback signals to EVC along the dorsal visual hierarchy to sustain subjective visual experience during WM maintenance.

### Differences between groups cannot be explained by univariate BOLD difference or eye movement patterns

Finally, we conducted several control analyses to ensure that the observed differences between groups were specific to differences in multivariate representational patterns. First, we tested whether these results could be attributed to variations in overall BOLD activity. We found no significant differences in BOLD activity within the EVC or IPS between groups in any task (Figure S9A-D).

Second, we examined whether eye movement patterns contributed to the observed neural differences. We analyzed eye gaze positions extracted via DeepMReye [58] during fMRI scanning. SVM decoding showed that gaze positions in neither group reliably predicted motion paths (Figure S9E-F). Additionally, we collected eye-tracking data during a behavioral session outside the scanner. Overall, decoding of motion paths from eye position data was weak in both typical imagers and aphantasics, and was only detectable from scattered time points during the perception task (Figure S9G-H). Again, the patterns of results did not align with the neural findings reported earlier. These findings together indicate that the observed differences between aphantasics and typical imagers cannot be explained by variations in overall BOLD activity or eye movement patterns.

## Discussion

We investigated the neural correlates of subjective visual experience during WM maintenance by comparing neural activity in typical imagers and aphantasics, who subjectively report no conscious visual experience during WM. Our findings provide converging evidence that feedback along a cortical hierarchy from IPS to EVC plays a critical role in supporting subjective visual experience during WM (Figure 6). First, aphantasics exhibited normal feedforward representations but weakened and unstable feedback representations of mnemonic stimuli in EVC compared to typical imagers. Second, IPS likely provided feedback signals necessary for normal visual imagery. Aphantasics showed reduced imagery generation signals in IPS, accompanied by reduced functional connectivity between EVC and IPS. Lastly, area V3AB served as a bottleneck along this pathway for the transition from visual to mnemonic codes. Notably, while the contralateral IPS was dysfunctional in aphantasia, ipsilateral IPS retained normal delay-period representations, which may underlie preserved WM abilities in aphantasia. Combined, these results indicate that feedback along a cortical hierarchy from IPS to EVC during WM, rather than activity within a single brain region, is critical for the conscious mental visualization of memory content during WM maintenance.

**Figure 6.**
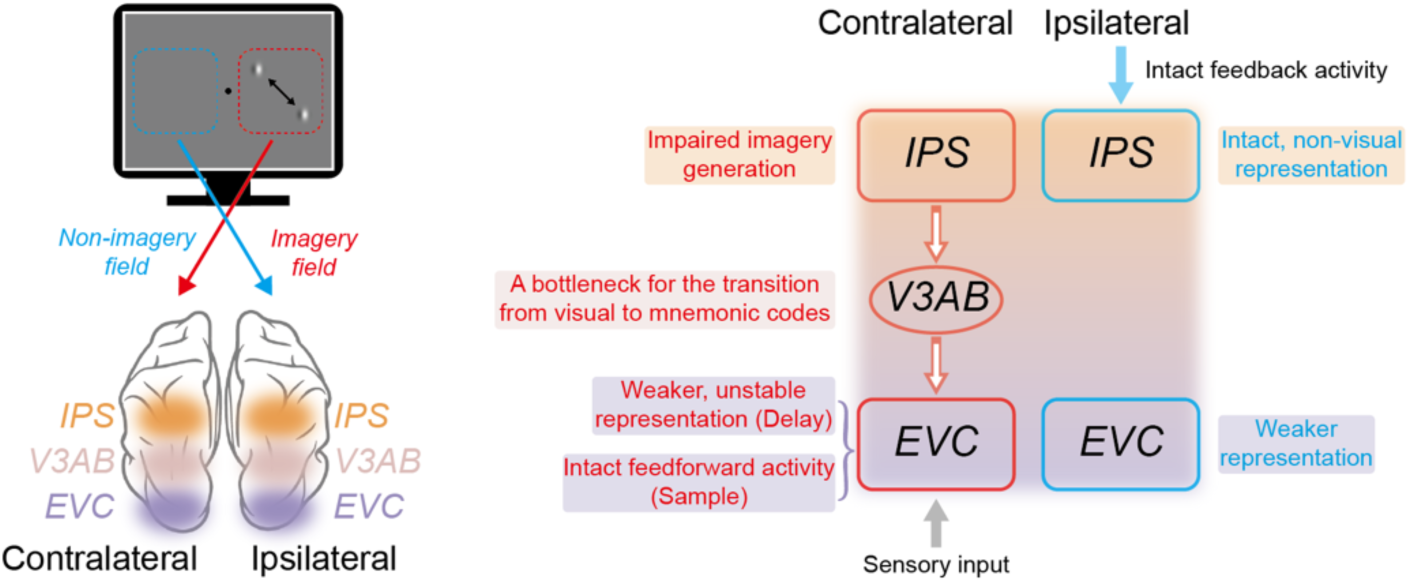
Schematic summary of main findings. During WM tasks, aphantasics differ from typical imagers in several key aspects: While the contralateral EVC represents feedforward sensory input normally, its WM representation becomes weak and unstable during the delay period. The contralateral IPS exhibits impaired imagery generation signals and reduced functional connectivity with contralateral EVC. In contrast, the ipsilateral IPS maintains intact, non-visual WM representation, which likely supports successful task performance independent of the impaired representation in ipsilateral EVC. Red and blue indicate regions contralateral and ipsilateral to the imagery field, respectively. Hollow and solid arrows denote impaired and intact functional connectivity, respectively.

### Feedback signals to EVC and subjective visual experience

Previous studies on the role of EVC in WM have largely focused on objective behavioral measures of WM such as precision and bias. Along this line of research, neural activity and stimulus representations in EVC have been shown to track WM precision [11,12] and to reflect response bias and drift over time [13,14,59]. Disrupting EVC activity also has a causal impact on WM fidelity [60]. These results together highlight the role of EVC in supporting WM maintenance. However, the necessity of EVC for WM has been debated, as electrophysiological studies in non-human primates often fail to find strong evidence for EVC involvement [61], but see [62]. By investigating a previously underexplored aspect of WM, the subjective visual experience, here we provide a unified account of EVC function. We propose that delay-period stimulus representations in EVC, which are considered to reflect feedback signals from higher-order cortices, are critical for subjective visual experience during WM maintenance, that is, conscious visualization of mnemonic content. Such visualization likely helps maintain visual details, thereby improving WM precision and performance. Conversely, impaired feedback to EVC leads to weakened stimulus representations, and consequently diminished conscious visual experience during WM.

WM is not a static storage but rather a dynamic, recoding process that serves subsequent task demands [28,53,54]. We further show that previously identified neural signatures of dynamic WM were also reduced with impaired feedback. These include the temporal stability of stimulus representations during WM delay, shared neural codes across WM tasks, and the dynamic transition from visual to mnemonic codes. In other words, impaired feedback to EVC not only precludes the transformation of stimulus representations into a memory format, but also disrupts the stable maintenance of these representations over time. These multifaceted roles of feedback signals may be crucial for shaping EVC representations to generate a coherent conscious experience.

### Impaired feedback develops along a cortical hierarchy between EVC and IPS

If feedback to EVC is crucial for subjective visual experience, where does this feedback signal originate? Previous work from our lab demonstrated imagery-dominant signals in IPS that are important for voluntary imagery generation in typical imagers [28]. This imagery dominance in IPS, however, was absent in aphantasics. In other words, neural signals for initiating imagery content were already impaired in aphantasics. Moreover, functional connectivity between EVC and IPS was also reduced in aphantasia. These results may explain why contralateral IPS also showed weakened stimulus representations towards later delay during illusion trials, because such voluntary control signals are critical for sustaining WM in the absence of external input. Importantly, however, impaired feedback to EVC as observed in our study cannot be fully explained by impaired imagery generation signals in IPS, because group differences were weak in IPS during perception trials, and group differences during illusion trials emerged later than those in EVC. These results suggest that impaired feedback to EVC can occur independent of IPS signals. Thus, impaired feedback affects EVC representations, and impaired imagery generation signals in IPS further reduces this feedback when visual imagery needs to be sustained over the memory delay. These processes may together contribute to diminished conscious visual experience in aphantasia.

Our finding that imagery-relevant signals gradually developed along the posterior-to-anterior visual hierarchy further supports a hierarchical model for subjective visual experience. The region V3AB, which lies between EVC and IPS, has been found to link perceptual and mnemonic processes during WM [28,63,64]. In our study, V3AB served as a transition point from sensory-driven signals to imagery-dominant representations during the sample period. During WM delay, it marked the transition from sensory-based to memory-based representations. Critically, in aphantasia, these transitions were blocked around V3AB, indicating a bottleneck that potentially impedes the propagation of feedback signals from IPS to EVC. These results indicate that EVC and IPS dysfunctions are not isolated, but instead exist along a continuum within the visual hierarchy, allowing impairments to emerge gradually at the network level.

It is noteworthy that, although the current study highlights IPS as a key source region of feedback signals, other studies on aphantasia have implicated additional regions in conveying feedback information across different cognitive processes, such as the hippocampus for autobiographical memory [65], the auditory cortex for cross-modal interactions [66], the orbitofrontal cortex for object imagery [51], and the prefrontal cortex during resting state [67]. This suggests that the source of feedback signals likely depends on the specific cognitive process. Our task required precise WM at specific retinotopic locations, making IPS particularly important due to its established roles in WM, mental imagery, and spatial cognition [4,6,24,28,40,68]. Moreover, feedback signals we observed in the IPS showed precise control over specific WM content at specific spatial locations, and therefore can be distinguished from content-invariant signals for more general control purposes. Future work could investigate the interplay between content-specific feedback signals in the IPS and general control signals from higher-order cortex, likely the prefrontal regions.

### Aphantasia involves a general deficit in feedback regardless of voluntary control

The multi-task design of the current study allowed us to examine whether impaired feedback to EVC in aphantasia reflects a selective impairment in voluntary imagery control or a more general deficit. Results from the illusion task support the latter. In illusion trials where minimal voluntary control was required during the sample period (i.e., involuntary imagery), aphantasics exhibited weakened stimulus representations in contralateral EVC, when no significant group differences were observed in contralateral IPS. Because the double-drift motion illusion is thought to recruit PFC [47], this result suggests that feedback to EVC (possibly from PFC) was already impaired during the sample period, even when participants could still consciously perceive the illusory stimuli. Therefore, aphantasics’ impairment in feedback to EVC is neither specific to the delay period nor to voluntary imagery control, and can occur even when stimulus representations in higher-order cortices are normal. This result has important implications for the characterization of aphantasia [69], indicating that aphantasia cannot be characterized simply as a deficit in voluntary imagery, but rather as a general deficit in feedback activity.

A related question, then, is why aphantasics can still experience the directional motion illusion during the sample period despite impaired feedback. We propose that continuous visual input from the moving Gabor, combined with its internal drift, enables higher-order regions to generate a directional bias. This bias is then integrated with sensory signals from EVC to generate conscious illusory perception, even without sufficient feedback to EVC during the sample period. Notably, this observation has important implications for theories of consciousness. EVC activity may not always reflect true perception, as perceptual experience can deviate from EVC signals during illusions [47]. Nevertheless, the presence of sensory signals in EVC is crucial for their integration with higher-order signals to form a coherent, conscious visual experience.

### Ipsilateral IPS supports successful WM performance without subjective visual experience

Despite these neural impairments, aphantasics were able to perform WM at a reasonable level. Consistent with this behavioral result, ipsilateral IPS contained stimulus representations that remained comparable between typical imagers and aphantasics. This suggests that ipsilateral IPS likely represents WM content in a compensatory, non-visual format to support behavior. Therefore, WM maintenance per se and its associated subjective conscious experience can be dissociated neurally. One remaining question is the origin of ipsilateral IPS signals. One possibility is that stimulus-specific signals in ipsilateral IPS arise from higher frontal regions [35,70]. Specifically, contralateral EVC in aphantasics encodes sensory-driven information normally and transmits it via feedforward pathways to higher-order regions, which then relay non-visual stimulus representations to ipsilateral IPS. This hypothesis, however, remains to be tested in future research.

## Conclusion

In summary, using aphantasia as a model, we demonstrate that normal, delay-period stimulus representations in the EVC depend on top-down feedback from higher-order IPS. Such feedback along a cortical hierarchy from IPS to EVC is critical for the subjective visual experience of visual content during WM maintenance, and impairments in this feedback pathway lead to deficits in visual consciousness in aphantasia.

## Methods

### Participants

A total of 54 participants were recruited for the study. Twenty of them were aphantasic participants, and the remaining participants were typical imagers. For aphantasics, two participants were excluded due to excessive head motion during fMRI scanning, and one participant withdrew midway, leaving 17 participants (5 males, mean age = 24.2 ± 3.5 years) in the fMRI session. Among these, seven participated in a second fMRI session. One was excluded due to excessive head motion, resulting in 6 participants (2 males, mean age = 23.0 ± 2.2 years) in the second session. All 17 participants additionally completed a behavioral session with eye-tracking. One participant was excluded due to technical issues during recording, leaving 16 participants in this session (6 males, mean age = 23.5 ± 3.5 years).

For typical imagers, fMRI data from 17 participants (6 males, mean age = 23.4 ± 1.5 years) were collected in a previous study from our lab [28]. Seventeen newly recruited participants completed the behavioral session with eye-tracking. One participant was excluded due to technical issues during recording, leaving 16 participants (6 males, mean age = 23.5 ± 3.5 years).

Sixteen aphantasic participants were recruited through social media platforms. The remaining 4 aphantasics and all typical imagers were recruited from the Chinese Academy of Sciences Shanghai Branch and surrounding areas. Aphantasia was further confirmed by a series of in-house questionnaires and interviews (detailed below).

All participants reported normal or corrected-to-normal vision and no history of neurological or psychiatric disorders. They provided written informed consent in accordance with protocols approved by the ethics committee of the Center for Excellence in Brain Science and Intelligence Technology, Chinese Academy of Sciences (CEBSIT-2020028), and received monetary compensation for their participation.

### Questionnaires and interviews for aphantasia

To assess participants’ self-assessed visual imagery abilities, both aphantasics and typical imagers completed the Vividness of Visual Imagery Questionnaire (VVIQ) [71] prior to the experiment. In the VVIQ, participants were instructed to mentally visualize 16 different scenes and then rate the vividness of the mental images they produced. Ratings were made on a 5-point scale from 1 (“No image at all; you only ‘know’ you are thinking of an object”) to 5 (“Perfectly clear and as vivid as normal vision”), with higher scores indicating stronger visual imagery ability.

In addition to the VVIQ, aphantasics completed a series of questionnaires to further confirm their absence of visual imagery. To characterize the imagery experience participants experienced, they were asked to imagine a yellow banana and select the picture that best matched their internal image (Figure S1B). Furthermore, six items from the Spontaneous Use of Imagery Scale [72] were included to assess participants’ spontaneous use of visual imagery in everyday situations (Figure S1C). Participants responded on a 5-point scale, where 1 indicated “Never appropriate” and 5 indicated “Completely appropriate.” To assess imagery abilities of other sensory modalities, nine items from the Plymouth Sensory Imagery Questionnaire [73] were selected - three items each targeting auditory, gustatory, and tactile imagery. Participants rated their imagery experience using a 6-point scale: where “1” indicated “Completely unable to imagine; only the name comes to mind,” “5” indicated “Image as clear and vivid as real life,” and “6” indicated “Unable to imagine because I have never encountered the described object or experience.”

Besides the questionnaires, each aphantasic participant completed a 60- to 80-minute interview, either before (8 participants) or after (9 participants) the experiment. They were asked a series of questions, including (1) their ability to generate imagery in other sensory modalities (i.e., auditory, olfactory, tactile, and gustatory imagery), (2) whether aphantasia affected their memory abilities, social interactions, face recognition, or emotional experiences, and (3) their dreaming experiences, including whether they experienced normal visual images during dreams.

### Experimental design and procedure

#### Apparatus and stimuli

All experimental stimuli were generated using MATLAB (The MathWorks) and Psychtoolbox-3. The screen background was gray (RGB: [128, 128, 128]) throughout the experiment. During behavioral training, visual stimuli were displayed on a HIKVISION LCD monitor (48 × 26.8 cm, 1920 × 1080 pixels at 60 Hz) with a viewing distance of 77 cm. Participants completed the task using a keyboard. During fMRI, visual stimuli were presented on a monitor positioned behind the scanner bore. For typical imagers, a SINORAD monitor screen was used (37.5 × 30 cm, 1280 × 1024 pixels, 60 Hz refresh rate). For individuals with aphantasia, a different monitor was employed (70 × 40 cm, 1920 × 1080 pixels, 120 Hz refresh rate). In both cases, the screen content was projected onto the participant’s field of view via a mirror mounted on the head coil. The viewing distance was 90.5 cm for typical imagers and 161 cm for aphantasics. Participants lay supine inside the scanner and performed the task using two SINORAD two-key button boxes.

#### Experimental procedure

The experiment consisted of a behavioral training session and an fMRI session, with an illusion size measurement test and three WM tasks in each session. A subset of aphantasic participants completed a second fMRI session. The only difference between the two fMRI sessions was whether stimuli appeared in the right (Session 1) or left (Session 2) visual field. All the experimental descriptions below were based on settings in Session 1. Throughout the experiment, participants were instructed to maintain fixation on a fixation point.

#### Illusion size measurement test

Prior to the WM tasks, participants completed a double-drift illusion measurement test following the procedure described in previous work [28]. Each trial began with the presentation of a Gabor pattern in the right peripheral visual field at an eccentricity of 8°. The stimulus consisted of a full-contrast sinusoidal grating windowed by a Gaussian envelope, with a spatial frequency of 0.94 cycles per degree and a diameter of 1.6° of visual angle. The Gabor pattern moved vertically at 5°/s for 2 s: first upward for 1 s, then downward for another 1 s. Concurrently, its internal texture either drifted leftward or rightward at a temporal frequency of 4 cycles per second or remained static (in the no-drift trials). The internal drift induced an illusory percept of left- or right-tilted motion, corresponding to the direction of the internal drift. Following stimulus presentation, a needle (0.05° wide, 5° long) appeared at fixation. Participants adjusted the needle’s orientation using four keys (two per hand) to align it with their perceived motion trajectory of the Gabor pattern. No time limit was imposed on responses. The initial orientation of the needle was randomized in each trial. The right-hand keys rotated the needle clockwisely (right key) or counterclockwisely (left key) in 1° increments per press, while the left-hand keys enabled 90° rapid rotation (left key) and response confirmation (right key).

Each run consisted of 30 trials, with 10 trials per stimulus, presented in a random order. Participants completed 3 to 4 runs in behavioral training and 2 to 3 runs inside the MRI scanner prior to the WM tasks. The illusion size measured in the final run was used as each participant’s individual illusion size in the subsequent WM tasks.

#### Working memory tasks

The WM tasks employed a delayed estimation trial structure with three variants: perception-based, imagery-based, and illusion-based WM (Figure 1A). The three tasks differed only in how the WM sample was delivered to participants.

In the perception task, a Gabor pattern (with its internal texture remaining static) moved back and forth along a tilted path in the right peripheral visual field at an eccentricity of 8°. The motion direction reversed every second; for example, for the left-tilted path, the Gabor first moved upward-left for 1 s, then reversed direction to move downward-right for another 1 s. The tilt angle of the motion path matched each participant’s measured illusion size. The vertical speed was fixed at 5°/s, while the horizontal speed and path length varied across participants depending on the path orientation.

In the imagery task, either the letter “L” or “R” was displayed at fixation as a symbolic cue, indicating whether participants should imagine a Gabor moving leftward or rightward during the subsequent delay period. The direction of the motion path for the imagined Gabor was required to match each participant’s perceived path direction during the perception or illusion tasks. No physical Gabor was presented in this task.

In the illusion task, the sample presentation was identical to the illusion size measurement test. The Gabor pattern moved vertically at 5°/s for 2 s (upward for 1 s, then downward for 1 s) in the peripheral visual field, while its internal texture drifted either leftward or rightward at a temporal frequency of 4 cycles per second (left- or right-tilted trials).

The sample period lasted 2 s in all tasks, followed by an 8.5-s delay period. During the delay, participants were instructed to mentally visualize the scene in which the perceived or imagined Gabor moved along the same motion path at the same peripheral location as in the perception or illusion trials. Following the delay, a needle appeared at fixation. Participants adjusted the needle’s orientation using button presses to report the previously perceived or imagined motion direction as accurately as possible. The adjustment procedure was identical to that in the illusion size measurement test, with a maximum response time of 4.5 s. Each trial ended with an inter-trial interval (ITI) of 4.5, 6, or 7.5 s. Each run consisted of 18 trials, with task type (imagery, perception, and illusion) and stimulus directions (i.e., leftward or rightward trajectory) balanced and randomly ordered across trials.

In the fMRI session, all 17 aphantasic participants completed 14 runs. Among the 17 typical imagers, one participant completed 12 runs, while the remaining completed 14 runs. The experimental setup was largely consistent for aphantasic participants and typical imagers. The main difference was that, among the aphantasic participants, seven followed the exact procedure described in previous work [28], where a black fixation point (0.3° in diameter) was displayed 3° horizontally to the left of the screen center. For those among these 7 participants who also completed a second fMRI session, the fixation point was displayed 3° horizontally to the right of the screen center. For all other participants, the fixation point remained at the center of the screen.

In the behavioral WM tasks of aphantasics and behavior-only typical imagers, after reporting the Gabor’s motion direction on each trial, participants additionally rated the vividness of their mental imagery during the delay period. Vividness was rated on a 4-point scale using 4 separate response keys with a maximum response time of 1.6 s: where “1” indicated no visual imagery; “4” indicated highly vivid imagery “as if actually seeing.” In addition, participants familiarized with the tasks and response keys in the first run, while subsequent runs were used for eye-tracking data collection. To ensure sufficient eye-tracking data, all aphantasic participants completed 5-6 runs, yielding at least 4 valid eye-tracking runs. For the 16 behavior-only typical imagers, fifteen completed 7-9 runs to ensure at least 6 valid eye-tracking runs, with one participant completing only 5 runs. To maintain consistency across groups, only the first 4 valid eye-tracking runs were included in the analysis.

### Data preprocessing and analyses

#### Behavioral analyses

Calculation of individual illusion size involved two steps: first, the perceived path direction was defined as the absolute difference between the reported orientation and the vertical orientation; second, illusion size was defined as the difference between the mean perceived direction in the double-drift trials and that in the no-drift trials.

For the WM tasks, recall error was quantified by calculating the angular difference between the reported orientation and the measured illusion size during the fMRI session. Additionally, the standard deviation (SD) of the reported orientation relative to the vertical orientation was calculated for each task to assess recall variability. When calculating vividness as well as recall error and variability, all trials were included. Due to a coding error, the recall error and variability for 3 typical imagers were not recorded. The behavioral results reported here are therefore based on data from the remaining 14 typical imagers.

#### Eye-tracking data analyses

During the behavioral WM tasks of aphantasics and behavior-only typical imagers, eye gaze position and pupil diameter were recorded using a desk-mounted infrared eye-tracking system (EyeLink 1000 Plus, SR Research, Ontario, Canada) at a sampling rate of 1000 Hz. The eye-tracking data were preprocessed following established methods [74] and custom scripts. First, raw data were filtered using a Savitzky–Golay finite impulse response (FIR) smoothing filter. Eye blinks and other artifacts were detected and removed using a velocity-based algorithm and acceleration criteria. Missing values resulting from eye blinks were reconstructed using spline, linear, or nearest-neighbor interpolation, depending on the timing of the blink within the trial: nearest-neighbor interpolation was applied when blinks occurred at the beginning or end of a trial, whereas linear or spline interpolation was used for blinks occurring in the middle of a trial, depending on the availability of valid data points on either side of the missing segment. The data were then baseline-corrected using the average signal from the −400 to 0 ms window relative to sample onset and downsampled to 100 Hz.

#### Data acquisition

MRI data form aphantasics were acquired using a 3 Tesla Siemens MRI scanner equipped with a 32-channel head coil at the Brain Imaging Center of the Institute of Neuroscience, Center for Excellence in Brain Science and Intelligence Technology, Chinese Academy of Sciences. Data from aphantasic particpants were obtained using a Prisma scanner, whereas data from typical imagers were obtained using a Tim Trio scanner. High-resolution T1-weighted anatomical images were obtained using an MPRAGE sequence with the following parameters: repetition time (TR) = 2300 ms, echo time (TE) = 3 ms, flip angle = 9°, matrix size = 256 × 256, 192 sequential sagittal slices, and isotropic voxel size = 1 mm³. Whole-brain functional images were collected using a multiband 2D gradient-echo echo-planar imaging (MB2D GE-EPI) sequence with a multiband acceleration factor of 2, repetition time (TR) = 1500 ms, echo time (TE) = 30 ms, flip angle = 60°, matrix size = 74 × 74, 46 axial slices, and isotropic voxel size = 3 mm³.

#### fMRI preprocessing

The preprocessing of fMRI data was performed using AFNI (https://afni.nimh.nih.gov/) [75]. For each functional run, the first five volumes were discarded to allow for signal stabilization. The data from each session were then aligned to the last volume of the final run within the same session, followed by alignment with the T1-weighted anatomical image from the first session (except that one participant’s data were aligned to the anatomical image from the second session). Subsequent steps included motion correction and detrending using linear, quadratic, and cubic regression. Voxel-wise signal time series were z-scored on a run-by-run basis unless otherwise specified.

#### Gaze position during fMRI task

To account for potential influences of eye movements on our results, we processed the fMRI data using a pretrained model (datasets_1to5) within DeepMReye’s Streamlit-based application [58]. The model generated ten predicted gaze positions for each fMRI volume. For subsequent analyses, we used the median of these ten gaze positions per volume.

#### ROI definition and voxel selection

We used the probabilistic atlas developed by Wang et al [76] to define individual anatomical regions of interest (ROIs). Probabilistic atlas masks were first inversely transformed into each participant’s native structural space, including visual areas V1, V2, V3, V3AB, hMT, MST, IPS0-5, and FEF. Subsequently, within each hemisphere, V1, V2, and V3 were combined to create early visual cortex (EVC) ROIs, IPS0-5 were combined for intraparietal sulcus (IPS) ROIs, hMT and MST were combined for MT+ ROIs, and the FEF mask was used for superior precentral sulcus (sPCS) ROIs in the frontal cortex.

To define functional ROIs exhibiting task-relevant activity, we selected voxels within each anatomical ROI based on task-related activation. First, a conventional mass-univariate GLM analysis was performed using AFNI. The GLM model incorporated six nuisance regressors accounting for head motion parameters, along with three regressors of interest representing the sample, delay, and probe periods of the task. These regressors were modeled as boxcar functions with durations of 2 s, 8.5 s, and 4.5 s, respectively, and were convolved with a canonical hemodynamic response function (HRF). The sample contrast was used to define functional EVC ROIs, while the delay contrast was used to define the remaining ROIs. In each case, the top 500 voxels showing the strongest activation for the respective contrast within the anatomical ROI were selected.

In addition to the ROIs mentioned above, we defined two additional individual anatomical ROIs—the occipital fusiform gyrus and the temporal occipital fusiform cortex using FSL’s Harvard–Oxford cortical lateralized atlas [77]. Given their limited voxel counts, all voxels were used without selection based on GLM beta values. Remaining steps followed the same procedure as above.

#### Calculation of BOLD time course

To compute BOLD time courses within each ROI during WM tasks, for each trial, the BOLD activity at each time point was calculated as the signal change relative to the first time point of that trial. For each WM task, the BOLD signal change was then averaged across all voxels within the ROI and across all trials of that task.

#### Support vector machine decoding

To assess the strength of path direction representations (leftward vs. rightward) for perceived or imagined Gabors across ROIs, we conducted ROI-based multivariate decoding analyses for each participant. We employed a two-class support vector machine (SVM) classifier using MATLAB’s built-in functions (‘fitcecoc’ for training and ‘predict’ for testing). For each ROI, we trained the SVM classifier using the preprocessed BOLD data from each WM task and tested it on the same task using a leave-one-trial-out cross-validation procedure. Within each training fold, the two classes were balanced, and each test fold contained two trials (one per class). Decoding accuracy was calculated as the proportion of correctly predicted trials across all test folds. To minimize potential bias associated with specific data partitioning, we randomized the assignment of trials to folds and repeated the entire cross-validation procedure 10 times. Final decoding accuracy was obtained by averaging accuracies across all iterations. To better illustrate the results, decoding accuracy was further averaged across time points within a task period (4-5th TR for sample, 7-8th TR for delay). We also performed cross-temporal and cross-task generalization analyses to compare path direction representations across time points and WM tasks, using the same leave-one-trial-out procedure. In the cross-temporal generalization analysis, classifiers were trained at each time point and tested across all time points in each task. For cross-task generalization, classifiers were trained on the perception task and tested on all three tasks. All decoding procedures were performed separately for each time point and ROI.

For all decoding analyses, trials with incorrect directional responses (e.g., responding “left” on rightward trials or vice versa) were excluded. The remaining trials were further balanced between left and right categories within each task to prevent overfitting during classifier training. Decoding of eye-tracking and fMRI gaze position data followed the same procedure. Results based on eye-tracking data were further smoothed using a 100-ms sliding window.

#### Representation similarity analyses (RSA)

To examine how the six motion paths (3 tasks × 2 motion directions) were represented across ROIs, we constructed two hypothesized representation dissimilarity models (RDMs). The first was

a visual RDM to capture differences in visual input during the sample period (Figure 5A): dissimilarity between different tasks was set to 1, while dissimilarity between the two motion paths (left vs. right) within the same task was set to 0.5. The second was a memory RDM to capture differences in the mental representations maintained during the delay period, in this model, no difference was expected across tasks if participants maintained the same WM information. Accordingly, if the maintained motion direction was the same across tasks, dissimilarity was set to 0; if the directions differed, dissimilarity was set to 1. A modified version of the visual RDM was also tested to validate the robustness of the effect regardless of the specific model used. The second visual RDM included parametric modulation of stimulus similarity (Figure S6A). For the illusion and perception tasks, dissimilarity increases with the angular difference in the physical motion path of the Gabor pattern. Angular difference (2 × illusion size) is maximal between the two oblique motion directions in the perception task and is set to a dissimilarity of 0.5. Within the illusion task, the two motion directions share the same physical path, resulting in a dissimilarity of 0. Between the perception and illusion tasks, the angular difference is smaller (1 × illusion size), with a dissimilarity of 0.25. The imagery task is most dissimilar from the other two tasks, in which no Gabor pattern was presented; thus, the dissimilarity is set to 1. Within this task, the dissimilarity between the two motion paths is set to 0.125.

Neural RDMs were constructed using SVM decoding results. Specifically, we performed pairwise decoding using SVMs with a leave-one-run-out cross-validation procedure. For each ROI and each time point, a 6×6 neural RDM was constructed based on pairwise decoding accuracies. The decoding accuracies served as measures of dissimilarity between each pair of motion directions, and diagonal elements of the RDMs were discarded. To account for collinearity between the two model RDMs, we computed two sets of partial Spearman correlation coefficients: one between neural RDMs and the visual RDM while controlling for the effect of the memory RDM, and another between neural RDMs and the memory RDM while controlling for the visual RDM. The resulting correlation coefficients quantified the similarity between neural RDM and different model RDMs at each time point within each ROI. Following the decoding analyses, trials with incorrect directional responses were excluded.

In addition to SVM decoding, we repeated the RSA by calculating the Mahalanobis distance between pairs of motion paths to construct neural RDMs. For each time point within each ROI, the data for each motion direction were randomly split into two halves. Both the training and testing data were then mean-centered by subtracting the mean of all six motion directions. Subsequently, Mahalanobis distance was computed between every motion path in the training and testing data, using the covariance matrix derived from combined training and testing data. This procedure was repeated 100 times, and the resulting mean distances across repetitions constituted the neural RDM for that ROI and time point. The diagonal elements were excluded following SVM-decoding RSA.

#### Generalized psychophysiological interactions

To investigate the functional connectivity between the EVC and IPS and between the V3AB and IPS, we performed generalized psychophysiological interaction (gPPI) analyses [57] using AFNI. We used preprocessed BOLD data prior to motion correction and detrending, with data scaled within each run using the AFNI scale block. Contralateral and ipsilateral IPS (defined by functional masks) were used as seed ROIs. EVC and V3AB located in the same hemisphere as the seed ROI, defined using its functional mask, were used as target ROIs.

For each participant, the gPPI GLM included the following regressors: (1) task regressors in the conventional GLM, consisting of 9 conditions (3 tasks × 3 periods), modeled as boxcar functions and convolved with a canonical HRF; (2) six head motion parameters as nuisance regressors; (3) the physiological BOLD time series extracted from the seed region, defined as the residual signals after regressing out variance associated with task regressors; and (4) nine psychophysiological interaction regressors (3 tasks × 3 periods).

To examine differences in functional connectivity between seed and target ROIs within each period, we compared the beta values of the psychophysiological interaction regressors across different tasks. These beta values were averaged across all voxels within the target ROI.

#### Statistical analyses

Behavioral differences between tasks were assessed using paired two-sample, two-tailed t-tests in MATLAB (ttest). All p-values were corrected for multiple comparisons using the false discovery rate (FDR) method for this and all subsequent analyses unless specified. Behavioral differences between groups were assessed using independent two-sample, two-tailed t-tests in MATLAB (ttest2).

To test the statistical significance for BOLD activity, partial Spearman correlation coefficients from RSA, and within-task decoding accuracy (with chance level subtracted), the following permutation procedure was used: For each group and task, at each time point, each participant’s result value was randomly multiplied by either +1 or −1, and the sign-flipped values were averaged across participants. This procedure was repeated 10000 times to generate a null distribution. The true group-average value was then compared against this null distribution, and p-values were calculated as the proportion of absolute values in the null distribution exceeding the absolute true value. Comparisons between tasks within the same group followed a similar procedure, except that sign-flipping was based on within-participant difference between tasks.

To assess differences between groups, results from all participants were pooled and randomly reassigned into two groups matching the original sample sizes, and the between-group difference for each randomized partition was computed. This procedure was repeated 10000 times to generate a null distribution. All remaining steps followed a similar procedure as above.

For cross-temporal and cross-task generalization analyses as well as for eye-tracking decoding, we applied a cluster-based permutation test. At each data point, one-sample one-tailed t-tests (ttest in MATLAB) were used to assess whether decoding accuracy exceeded chance level (*p* < 0.05). Spatiotemporally (for cross-generalization) or temporally (for eye-tracking) adjacent significant data points were grouped into clusters, and the sum of t-values within each cluster was computed as the cluster statistic. A null distribution was generated by subtracting chance level from each participant’s decoding accuracy, randomly flipping the sign, and repeating the same t-test and clustering procedure. For each permutation, the largest cluster-level sum of t-values was recorded. This procedure was repeated 10000 times to construct the null distribution. Clusters were deemed significant if their sum of t-values exceeded the 99th percentile of the null distribution (*p* < 0.01).

## Acknowledgements

This work was supported by the Ministry of Science and Technology of China (STI2030-Major Projects 2021ZD0204202, 2021ZD0203701), the National Natural Science Foundation of China (32271089), CAS Project for Young Scientists in Basic Research (YSBR-071), and the Strategic Priority Research Program of the Chinese Academy of Sciences (XDB1010202) to Q.Y..

## Supplemental Information

**Figure S1.**
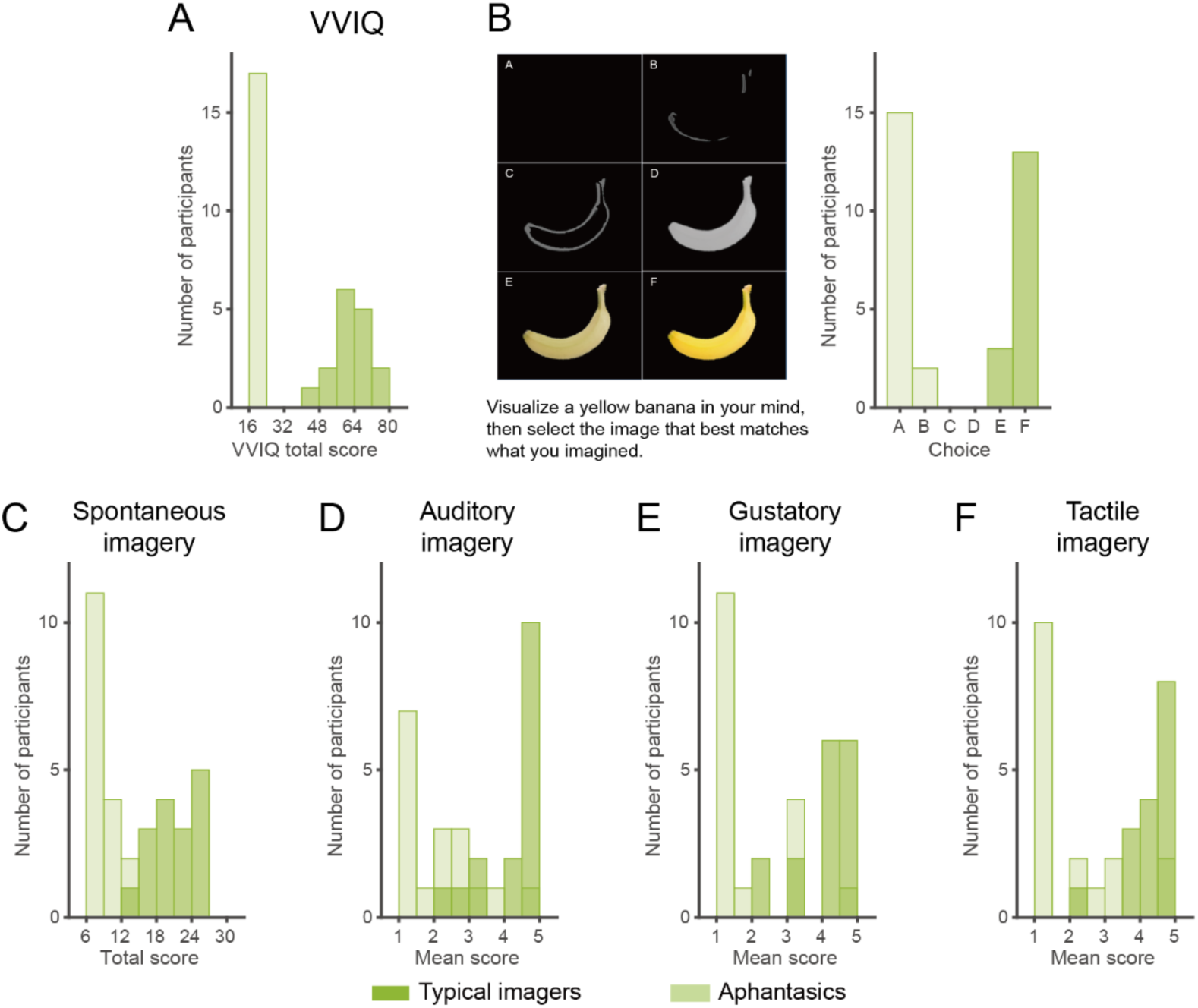
Summary of questionnaire scores for typical imagers and aphantasics. (A) Histogram of total VVIQ scores for typical imagers (dark green) and aphantasics (light green). (B) Imagery visualization. Left panel: example images corresponding to each choice. Right panel: frequency of choices for typical imagers and aphantasics. (C). Histogram of total scores on selected items from the Spontaneous Use of Imagery Scale for typical imagers and aphantasics. (D–F) Scores on the Plymouth Sensory Imagery Questionnaire subscales. Histograms of mean scores on the (D) auditory, (E) gustatory, and (F) tactile imagery subscales for typical imagers and aphantasics.

**Figure S2.**
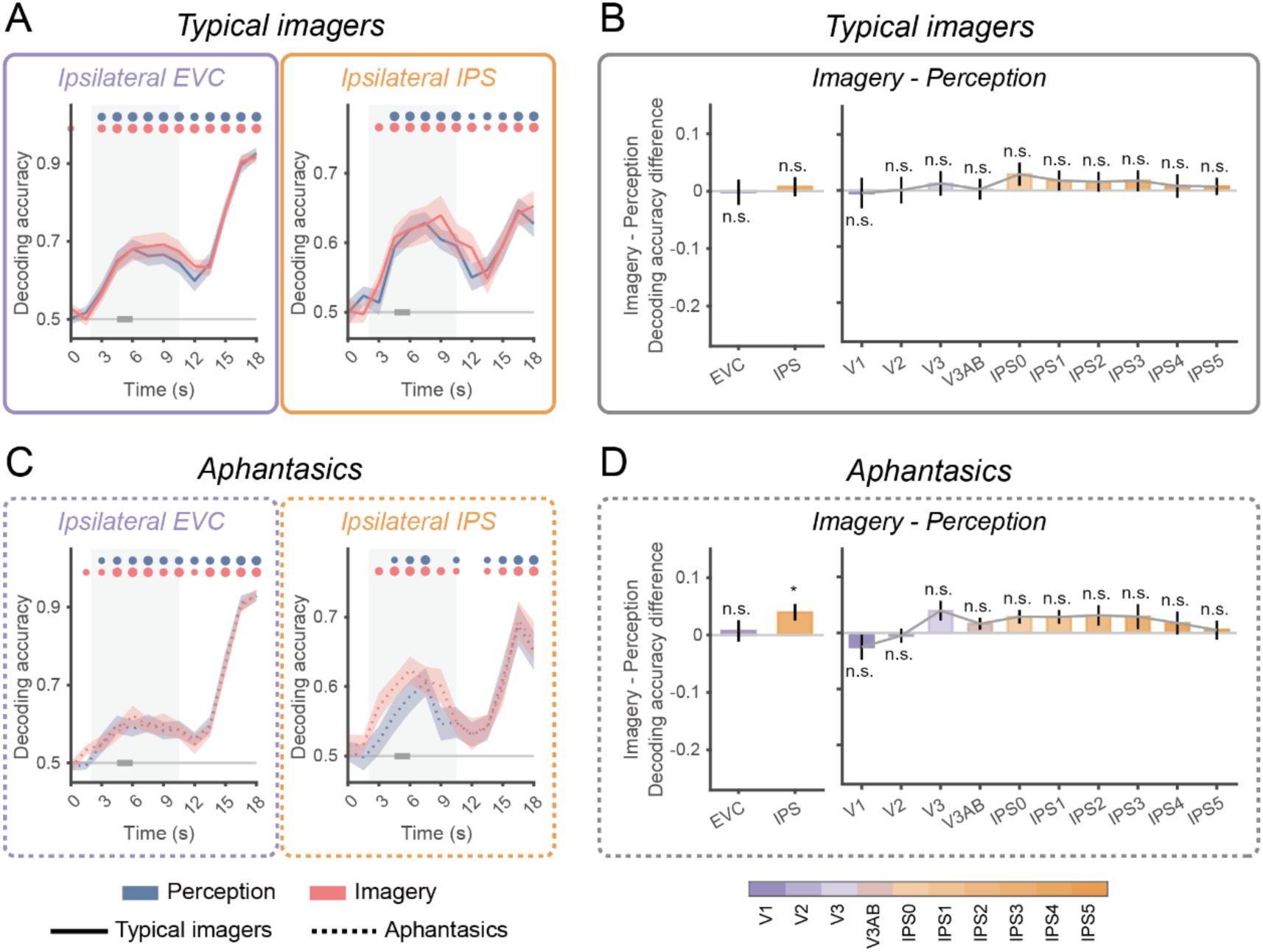
Imagery dominance in ipsilateral EVC and IPS for typical imagers and aphantasics. (A) Time course of decoding accuracy for perception (blue) and imagery (red) in ipsilateral EVC (left) and IPS (right). Circles on the top denote significant time points, and circle size reflects significance: large, *p* < 0.001; medium, *p* < 0.01; small, *p* < 0.05. All p-values were FDR-corrected. (B) Imagery dominance in typical imagers. Left panel: Imagery dominance in ipsilateral EVC and IPS during the sample period (time windows indicated by thick gray bars in A), defined as the difference in decoding accuracy between imagery and perception tasks. Error bars indicate ±1 SEM. Asterisks denote significant differences between perception and imagery within the corresponding ROI. Right panel: Imagery dominance across ipsilateral visual hierarchy (V1 to IPS5), using the same conventions. Significance levels: ***, *p* < 0.001; **, *p* < 0.01; *, *p* < 0.05; n.s., *p* ≥ 0.05. All p-values were FDR-corrected. (C) Same layout and conventions as (A), with results from aphantasics (dashed lines). (D) Same layout and conventions as (B), with results from aphantasics.

**Figure S3.**
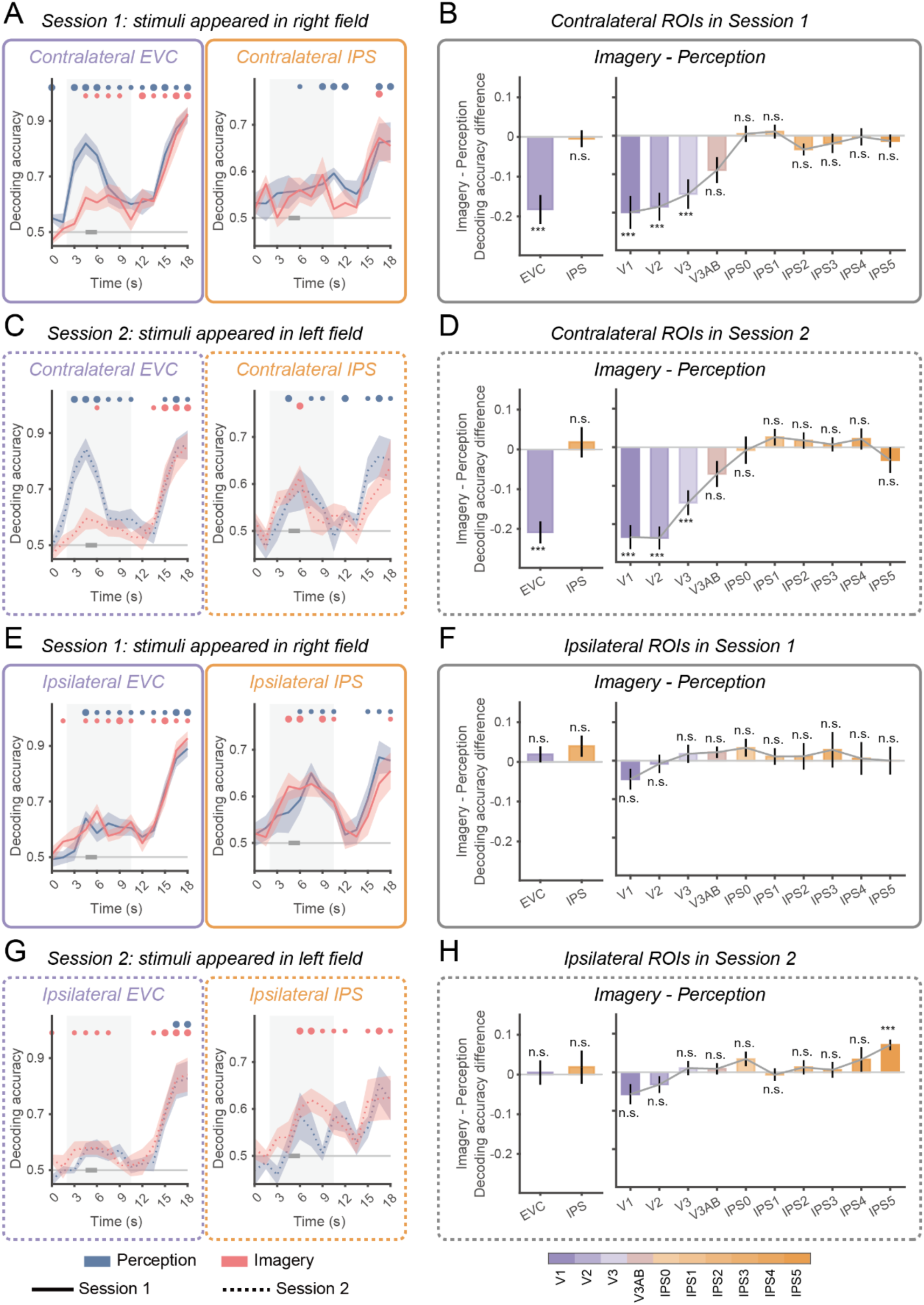
Comparison of imagery dominance between sessions (right versus left visual field) in EVC and IPS of aphantasics (n = 6). (A) Time course of decoding accuracy for perception (blue) and imagery (red) in contralateral EVC (left) and IPS (right), from Session 1 where stimuli appeared in the right visual field. Circles on the top denote significant time points, and circle size reflects significance: large, *p* < 0.001; medium, *p* < 0.01; small, *p* < 0.05. All p-values were FDR-corrected. (B) Imagery dominance in aphantasics from Session 1. Left panel: Imagery dominance in contralateral EVC and IPS during the sample period (time windows indicated by thick gray bars in A), defined as the difference in decoding accuracy between imagery and perception tasks. Error bars indicate ±1 SEM. Asterisks denote significant differences between perception and imagery within the corresponding ROI. Right panel: Imagery dominance across contralateral visual hierarchy (V1 to IPS5), using the same conventions. Significance levels: ***, *p* < 0.001; **, *p* < 0.01; *, *p* < 0.05; n.s., *p* ≥ 0.05. All p-values were FDR-corrected. (C) Same layout and conventions as (A), with results from Session 2 where stimuli appeared in the left visual field. (D) Imagery dominance in Session 2 (contralateral). Same layout and conventions as (B). (E) Same layout and conventions as (A), with results from ipsilateral ROIs in Session 1. (F) Imagery dominance in Session 1 (ipsilateral). Same layout and conventions as (B), (G) Same layout and conventions as (A), with results from ipsilateral ROIs in Session 2. (H) Imagery dominance in Session 2 (ipsilateral). Same layout and conventions as (B).

**Figure S4.**
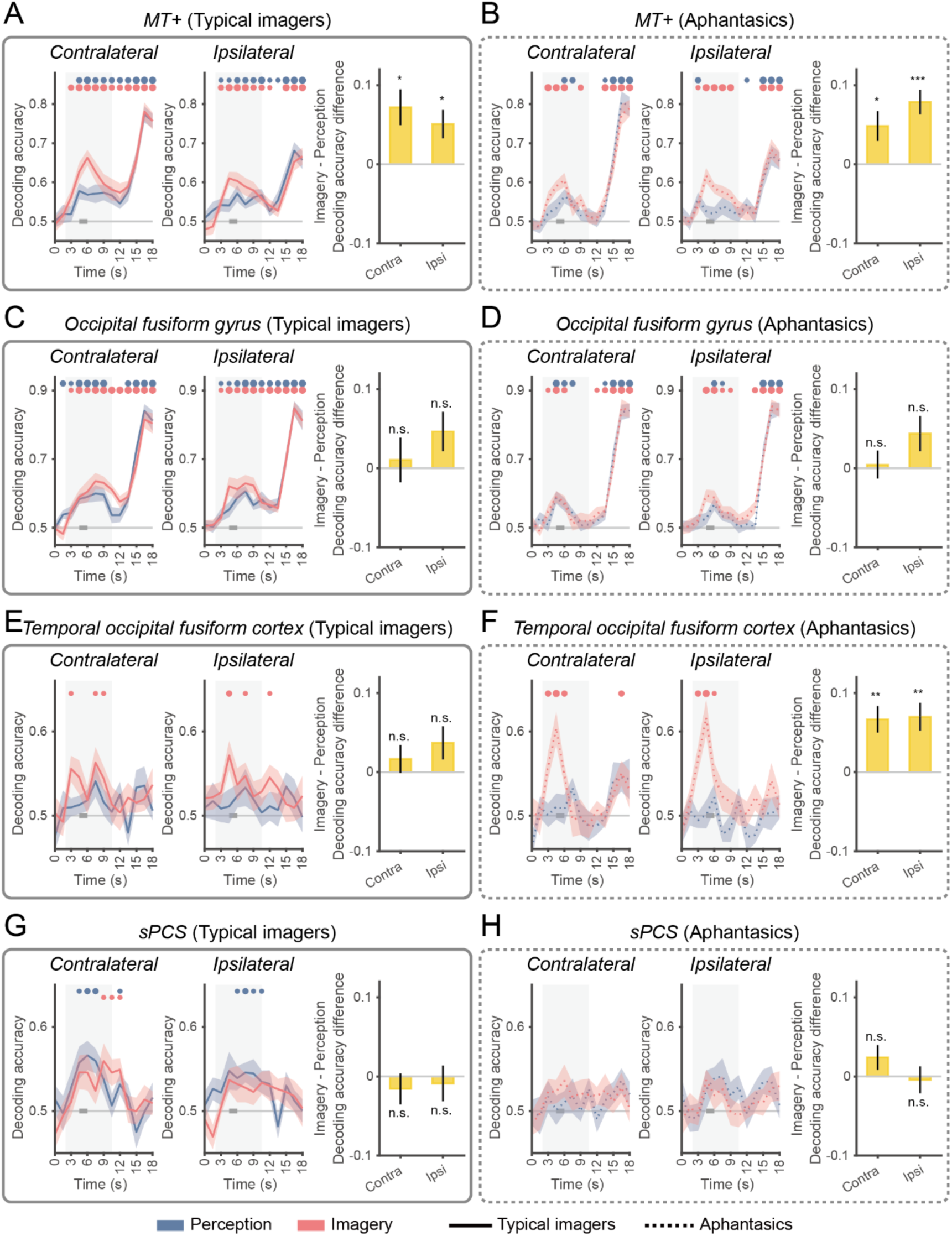
Imagery dominance in MT+, occipital fusiform gyrus, temporal occipital fusiform cortex and sPCS for typical imagers and aphantasics. (A) Typical imagers (MT+). Decoding accuracy for perception (blue) and imagery (red) in contralateral (left panel) and ipsilateral MT+ (middle panel). Significant time points are marked above. Circle size indicates significance (large: p<0.001; medium: p<0.01; small: p<0.05). Right panel: Imagery dominance in contralateral and ipsilateral MT+. Significance levels: ***p<0.001, **p<0.01, *p<0.05, n.s. p≥0.05. All p-values are FDR-corrected. (B–H) Results follow the identical layout as in (A), corresponding respectively to: (B) Aphantasics (MT+), (C) Typical imagers (occipital fusiform gyrus), (D) Aphantasics (occipital fusiform gyrus), (E) Typical imagers (temporal occipital fusiform cortex), (F) Aphantasics (temporal occipital fusiform cortex), (G) Typical imagers (sPCS), (H) Aphantasics (sPCS). Abbreviations: Contra, contralateral; Ipsi, ipsilateral.

**Figure S5.**
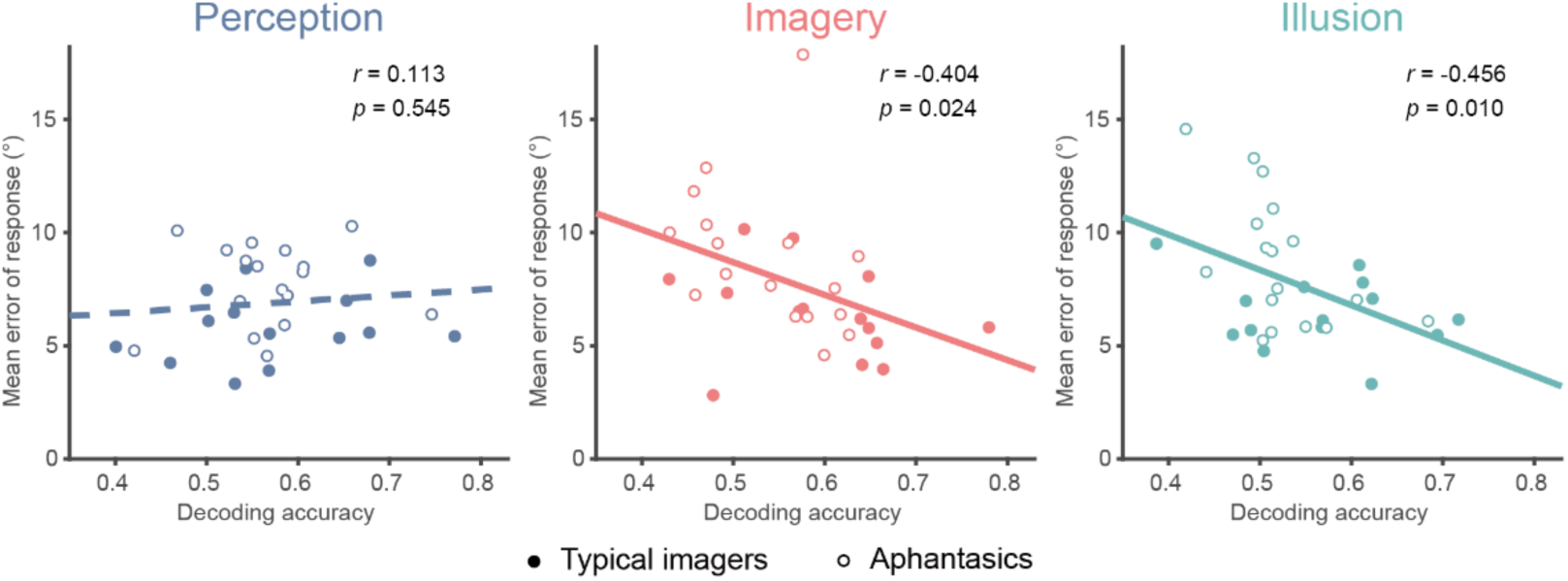
Correlation between IPS decoding accuracy and response error. Left panel: Each circle represents individual IPS decoding accuracy (during the delay period) and corresponding response error in the perception (left panel), imagery (middle panel), and illusion (right panel) tasks. Filled and hollow circles denote typical imagers and aphantasics, respectively. Solid and dashed lines denote significant (*p* < 0.05) and non–significant correlations, respectively.

**Figure S6.**
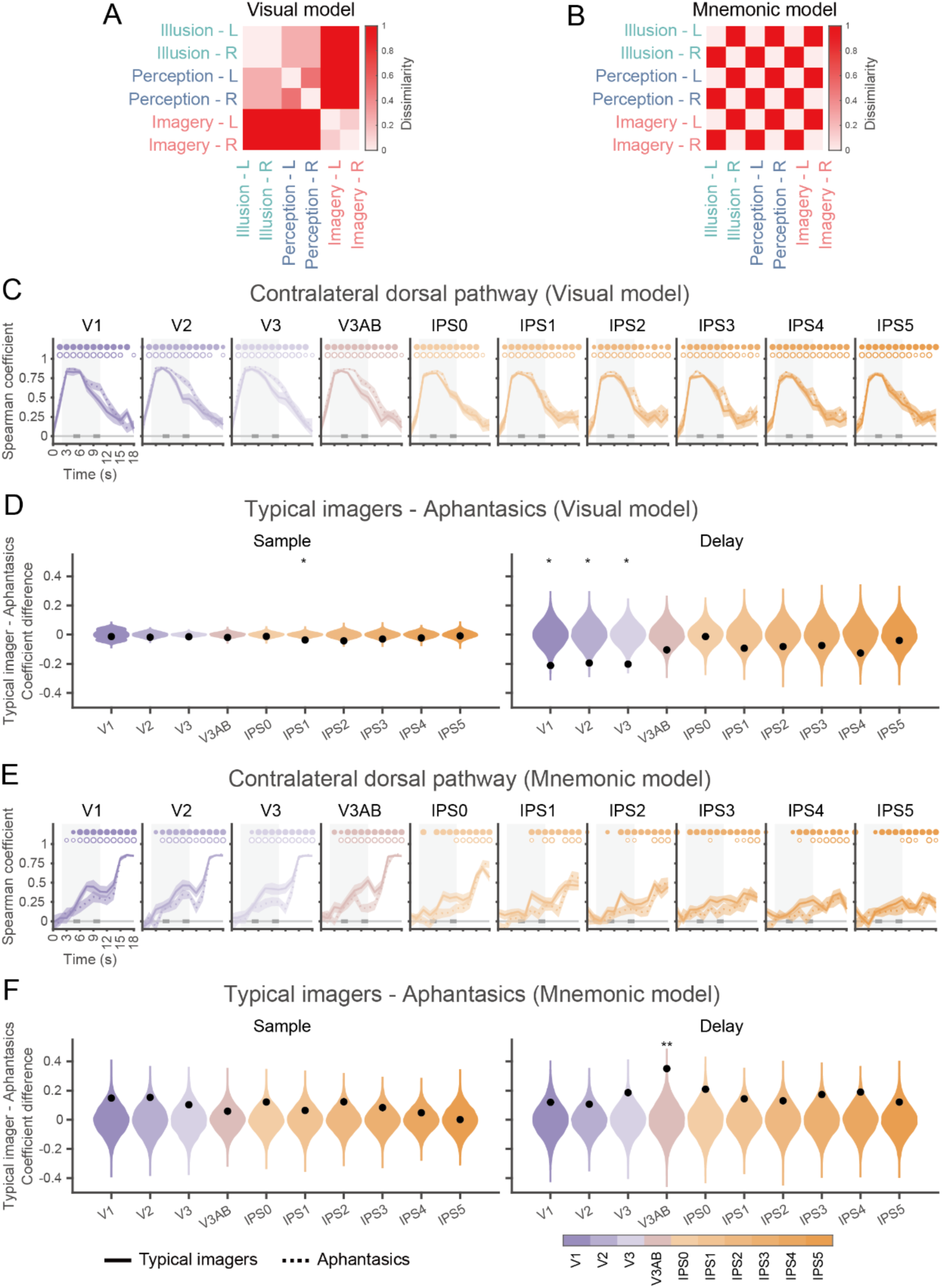
Temporal dynamics of motion path representational formats from contralateral V1 to IPS5 in typical imagers and aphantasics using an alternative visual model. (A) Visual model. Hypothetical dissimilarity structure among six motion paths (3 tasks × 2 paths) that reflects visual differences during the sample period. Dissimilarity between conditions is adjusted based on parametric modulation of visual presentation (see methods). (B) Mnemonic model. Hypothetical dissimilarity structure that reflects WM differences during the delay period, based solely on motion-path identity: dissimilarity is 0 for identical paths and 1 otherwise. (C) Correlation with the visual model. Time-resolved Spearman correlation between neural RDMs (contralateral V1 to IPS5, left to right) and the visual model. Solid line: typical imagers; dashed line: aphantasics. Significant time points are indicated above (filled circles: typical imagers; hollow circles: aphantasics). Circle size reflects significance: large, *p* < 0.001; medium, *p* < 0.01; small, *p* < 0.05. All p-values were FDR-corrected. (D) Difference in RDM similarity between typical imagers and aphantasics. Left panel: The violin plots depict the null distribution of group differences in Spearman correlation coefficients for the visual model during the sample period (left thick gray bar in C), derived from permutation testing. The black dots denote true group differences. Right panel: Corresponding results for the delay period (right thick gray bar in C). Asterisks denote statistical significance: *** *p* < 0.001; ** *p* < 0.01; * *p* < 0.05. All p-values were FDR-corrected. (E) Same layout and conventions as (C), but for mnemonic model. (F) Same layout and conventions as (D), but for mnemonic model.

**Figure S7.**
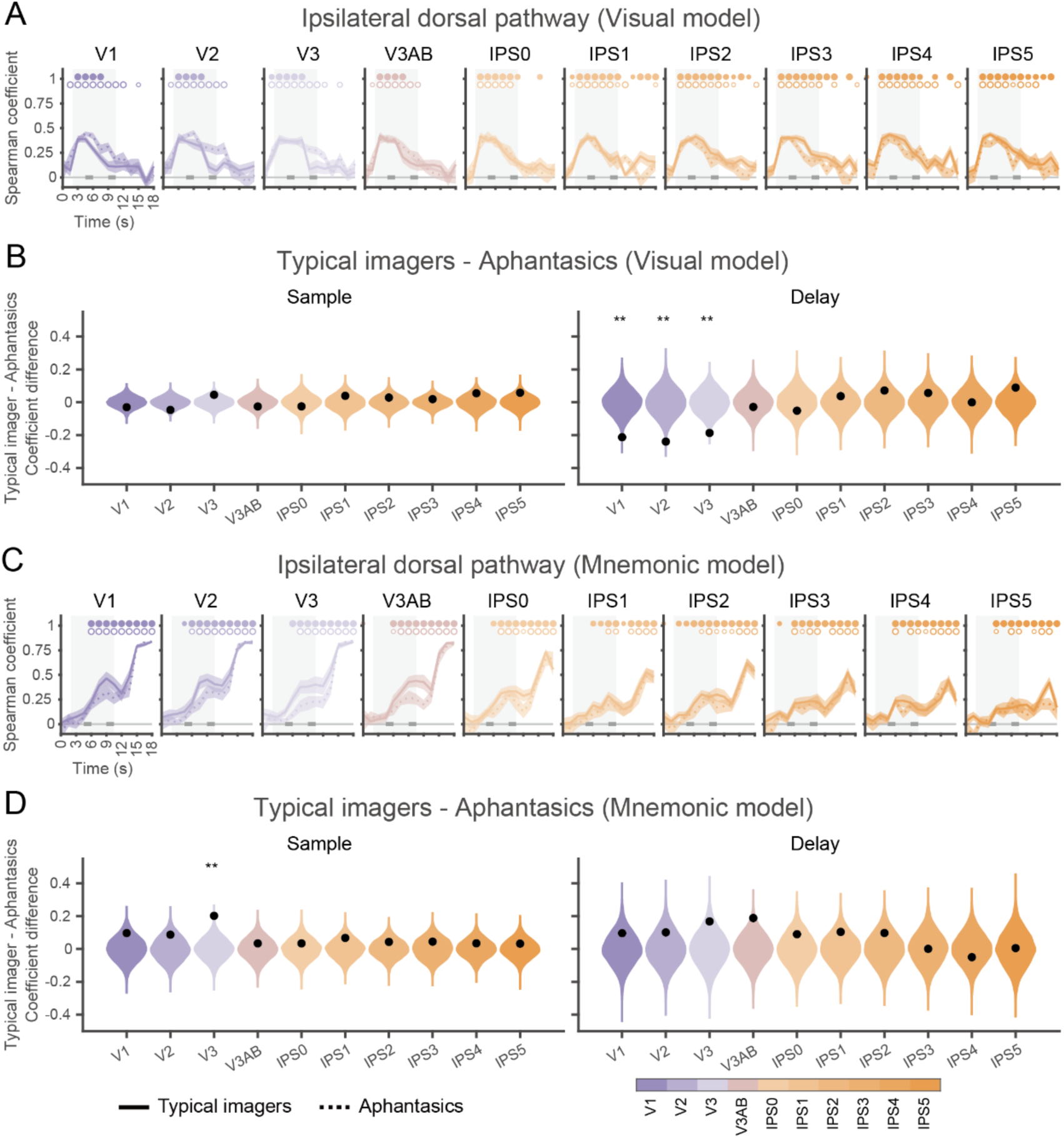
Temporal dynamics of motion path representational formats from ipsilateral V1 to IPS5 in typical imagers and aphantasics. (A) Similarity between neural RDMs and the visual model (in Figure 5A) over time. Spearman correlations between neural RDMs (ipsilateral V1 to IPS5, left to right) and the visual model for typical imagers (solid line) and aphantasics (dashed line). Significant time points are indicated above (filled circles: typical imagers; hollow circles: aphantasics). Circle size reflects significance: large, p < 0.001; medium, p < 0.01; small, p < 0.05. All p-values were FDR-corrected. (B) Difference in RDM similarity between typical imagers and aphantasics. Left panel: The violin plots depict the null distribution of group differences in Spearman correlation coefficients for the visual model during the sample period (left thick gray bar in A), derived from permutation testing. The black dots denote true group differences. Right panel: Corresponding results for the delay period (right thick gray bar in A). Asterisks denote statistical significance: *** *p* < 0.001; ** *p* < 0.01; * *p* < 0.05. All p-values were FDR-corrected. (C) Same layout and conventions as (A), but for the mnemonic model (in Figure 5B). (D) Same layout and conventions as (B), but for the mnemonic model.

**Figure S8.**
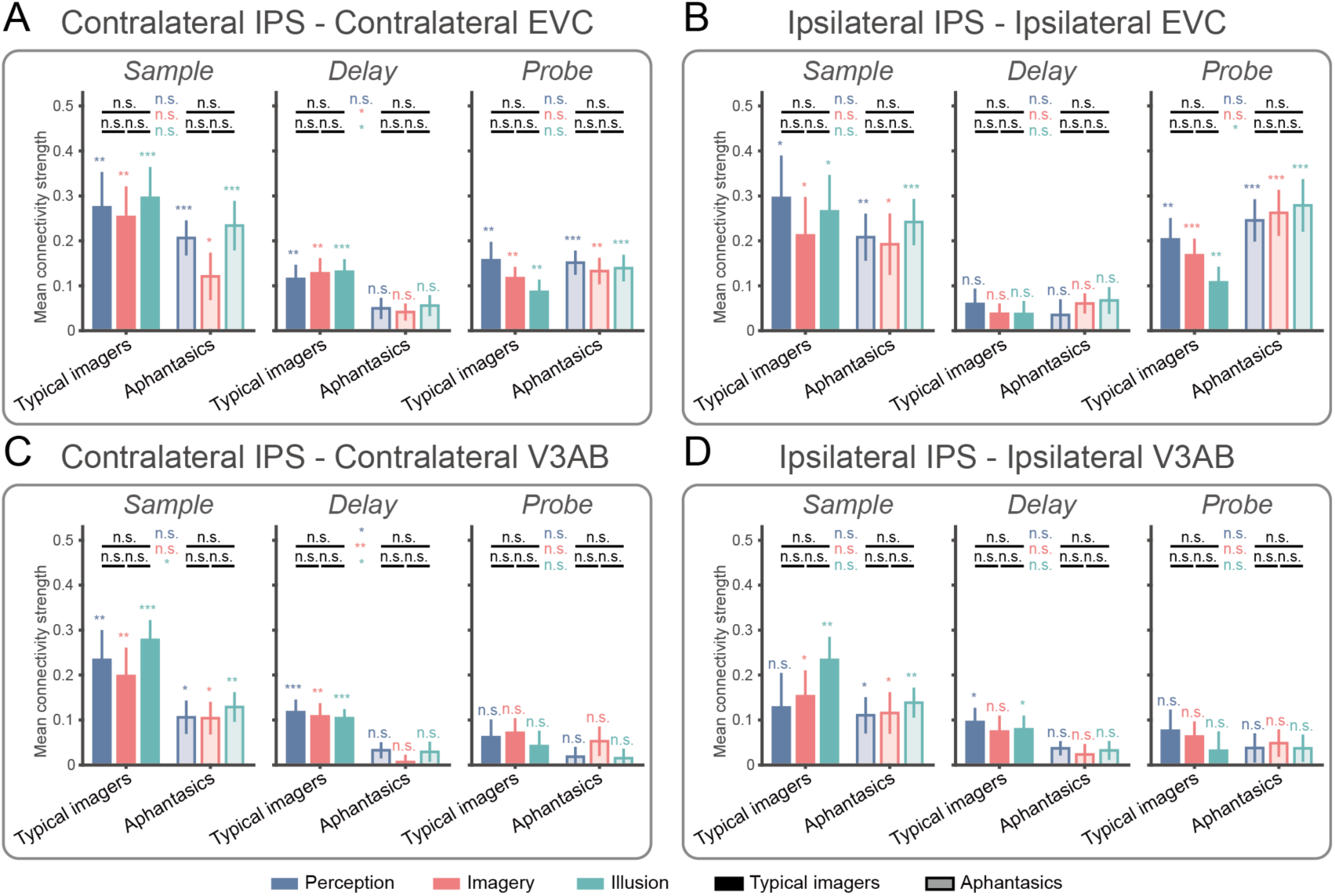
Functional connectivity between IPS and visual areas. (A) Functional connectivity between contralateral IPS (seed) and contralateral EVC for the sample (left), delay (middle), and probe (right) periods, calculated using gPPI method. Colored asterisks above individual error bars indicate significant connectivity within a specific group and task, black asterisks above bars denote significant differences between tasks within the same group, colored asterisks positioned between groups indicate significant differences between groups for the corresponding task. (B) Same layout and conventions as (A), with results from ipsilateral IPS (seed) and EVC. (C) Same layout and conventions as (A), with results from contralateral IPS (seed) and V3AB. (D) Same layout and conventions as (A), with results from ipsilateral IPS (seed) and V3AB. Significance levels: ***, *p* < 0.001; **, *p* < 0.01; *, *p* < 0.05; n.s., *p* ≥ 0.05. All p-values were FDR-corrected except those for between-group differences.

**Figure S9.**
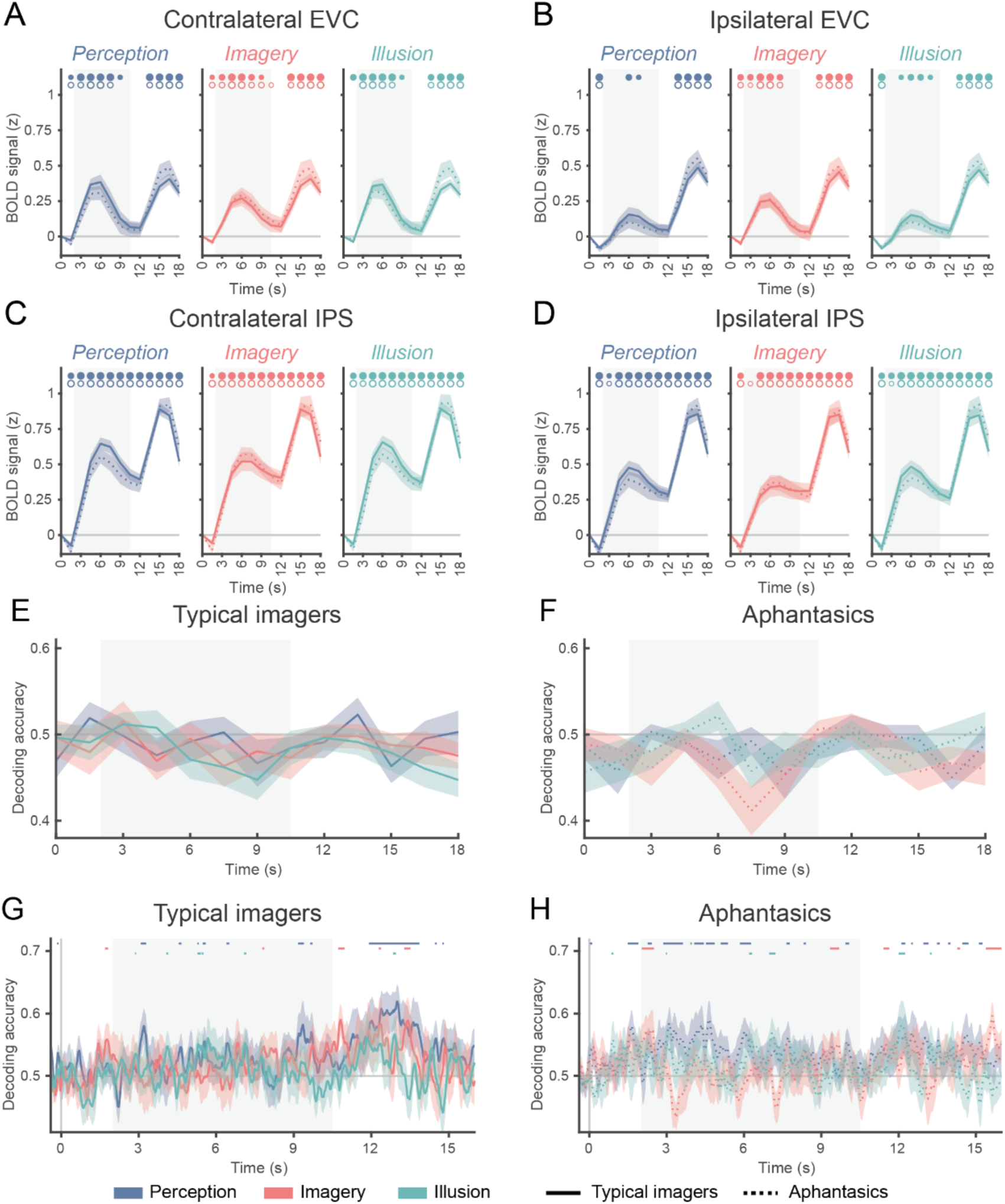
BOLD time course and decoding of eye–movement-related data for typical imagers and aphantasics. (A) BOLD time course for perception, imagery, and illusion tasks in contralateral EVC for typical imagers (solid line) and aphantasics (dashed line). Circles on the top denote significant time points (filled circles: typical imagers; hollow circles: aphantasics). Circle size reflects significance: large, *p* < 0.001; medium, *p* < 0.01; smalle, *p* < 0.05. All p-values were FDR-corrected. (B–D) Same layout and conventions as (A), with results from (B) ipsilateral EVC, (C) contralateral IPS, (D) ipsilateral IPS. (E) Time course of motion-path decoding from eye gaze position in typical imagers. Gaze position was extracted from EPI data using DeepMReye. (F) Same conventions as (E), but with results from aphantasics. (G) Time course of motion-path decoding from eye-tracking data in typical imagers. Eye-tracking data were collected during the behavioral session. Clusters with *p* < 0.05 after cluster-based correction are shown. (H) Same conventions as (G), but with results from aphantasics.

**Table S1.** Interview summary for aphantasics.

| Participant | Auditory<br>imagery | Olfactory<br>imagery | Gustatory<br>imagery | Tactile<br>imagery | Impact on<br>memory | Impact on<br>social<br>interactions | Impact on face<br>memory | Impact on<br>emotions | Normal images<br>during dreams |
| --- | --- | --- | --- | --- | --- | --- | --- | --- | --- |
| AA | No | No | No | No | No | No | No | No | Yes |
| AB | Yes | Yes | Yes | Yes | Yes | Yes | Yes | No | No |
| AC | No | No | No | No | No | No | No | No | No |
| AD | No | No | No | No | Yes | Yes | Yes | No | No |
| AE | Yes | Yes | Yes | Yes | No | No | Yes | No | Yes |
| AF | Yes | No | No | No | Yes | No | Yes | Yes | Not sure |
| AG | Yes | Yes | Yes | Yes | No | No | Yes | No | Yes |
| AH | No | No | No | No | No | No | No | Not sure | No |
| AI | Yes | Yes | Yes | Yes | Not sure | Yes | Yes | Yes | Not Sure |
| AJ | No | No | No | No | Yes | No | No | No | No |
| AK | Yes | Yes | Yes | Yes | No | No | No | No | Not sure |
| AL | No | No | No | No | No | No | Yes | No | No |
| AM | No | No | No | No | Yes | No | Yes | Not Sure | Yes |
| AN | No | No | No | No | Yes | No | Yes | Yes | No |
| AO | Yes | No | No | No | Yes | No | Yes | No | No |
| AP | No | No | No | No | No | No | Yes | No | Not Sure |
| AQ | No | No | No | No | Yes | Not Sure | Yes | No | Yes |
| **Summary** |  |  |  |  |  |  |  |  |  |
| (% of yes<br>response) | 7/17 | 5/17 | 5/17 | 5/17 | 8/17 | 3/17 | 12/17 | 3/17 | 5/17 |

## References

1. Baddeley A. Working memory: looking back and looking forward. Nat Rev Neurosci. 2003;4(10):829–39. doi: 10.1038/nrn1201.

2. Muraki EJ, Speed LJ, Pexman PM. Insights into embodied cognition and mental imagery from aphantasia. Nature Reviews Psychology. 2023;2(10):591–605. doi: 10.1038/s44159-023-00221-9.

3. Harrison SA, Tong F. Decoding reveals the contents of visual working memory in early visual areas. Nature. 2009;458(7238):632–5. doi: 10.1038/nature07832.

4. Ester EF, Sprague TC, Serences JT. Parietal and Frontal Cortex Encode Stimulus-Specific Mnemonic Representations during Visual Working Memory. Neuron. 2015;87(4):893–905. doi: 10.1016/j.neuron.2015.07.013.

5. Christophel TB, Klink PC, Spitzer B, Roelfsema PR, Haynes JD. The Distributed Nature of Working Memory. Trends Cogn Sci. 2017;21(2):111–24. doi: 10.1016/j.tics.2016.12.007.

6. Yu Q, Shim WM. Occipital, parietal, and frontal cortices selectively maintain task-relevant features of multi-feature objects in visual working memory. Neuroimage. 2017;157:97–107. doi: 10.1016/j.neuroimage.2017.05.055.

7. Shao Z, Zhang M, Yu Q. Stimulus representation in human frontal cortex supports flexible control in working memory. Elife. 2025;13:RP100287. doi: 10.7554/eLife.100287.

8. Lee SH, Kravitz DJ, Baker CI. Goal-dependent dissociation of visual and prefrontal cortices during working memory. Nat Neurosci. 2013;16(8):997–9. doi: 10.1038/nn.3452.

9. D’Esposito M, Postle BR. The cognitive neuroscience of working memory. Annu Rev Psychol. 2015;66:115–42. doi: 10.1146/annurev-psych-010814-015031.

10. Scimeca JM, Kiyonaga A, D’Esposito M. Reaffirming the Sensory Recruitment Account of Working Memory. Trends Cogn Sci. 2018;22(3):190–2. doi: 10.1016/j.tics.2017.12.007.

11. Emrich SM, Riggall AC, Larocque JJ, Postle BR. Distributed patterns of activity in sensory cortex reflect the precision of multiple items maintained in visual short-term memory. J Neurosci. 2013;33(15):6516–23. doi: 10.1523/JNEUROSCI.5732-12.2013.

12. Gosseries O, Yu Q, LaRocque JJ, Starrett MJ, Rose NS, Cowan N, et al. Parietal-Occipital Interactions Underlying Control- and Representation-Related Processes in Working Memory for Nonspatial Visual Features. J Neurosci. 2018;38(18):4357–66. doi: 10.1523/JNEUROSCI.2747-17.2018.

13. Hallenbeck GE, Sprague TC, Rahmati M, Sreenivasan KK, Curtis CE. Working memory representations in visual cortex mediate distraction effects. Nat Commun. 2021;12(1):4714. doi: 10.1038/s41467-021-24973-1.

14. Gu H, Lee J, Kim S, Lim J, Lee HJ, Lee H, et al. Attractor dynamics of working memory explain a concurrent evolution of stimulus-specific and decision-consistent biases in visual estimation. Neuron. 2025;113(20):3476–90 e9. doi: 10.1016/j.neuron.2025.07.003.

15. Lorenc ES, Sreenivasan KK, Nee DE, Vandenbroucke ARE, D’Esposito M. Flexible coding of visual working memory representations during distraction. J Neurosci. 2018. doi: 10.1523/JNEUROSCI.3061-17.2018.

16. Henderson MM, Rademaker RL, Serences JT. Flexible utilization of spatial- and motor-based codes for the storage of visuo-spatial information. Elife. 2022;11. doi: 10.7554/eLife.75688.

17. Zeman A, Dewar M, Della Sala S. Lives without imagery - Congenital aphantasia. Cortex. 2015;73:378–80. doi: 10.1016/j.cortex.2015.05.019.

18. Zeman A. Aphantasia and hyperphantasia: exploring imagery vividness extremes. Trends Cogn Sci. 2024;28(5):467–80. doi: 10.1016/j.tics.2024.02.007.

19. Keogh R, Wicken M, Pearson J. Visual working memory in aphantasia: Retained accuracy and capacity with a different strategy. Cortex. 2021;143:237–53. doi: 10.1016/j.cortex.2021.07.012.

20. Kay L, Keogh R, Pearson J. Slower but more accurate mental rotation performance in aphantasia linked to differences in cognitive strategies. Conscious Cogn. 2024;121:103694. doi: 10.1016/j.concog.2024.103694.

21. Knight KF, Milton F, Zeman A. Aphantasia and visual working memory: No direct evidence of impaired visual working memory in aphantasics, either in behavioral performance or the accuracy of a multivoxel pattern classifier. Neuropsychologia. 2026:109430. doi: 10.1016/j.neuropsychologia.2026.109430.

22. Cabbai G, Racey C, Simner J, Dance C, Ward J, Forster S. Sensory representations in primary visual cortex are not sufficient for subjective imagery. Curr Biol. 2024;34(21):5073–82 e5. doi: 10.1016/j.cub.2024.09.062.

23. Chang S, Zhang X, Cao Y, Pearson J, Meng M. Imageless imagery in aphantasia revealed by early visual cortex decoding. Curr Biol. 2025;35(3):591–9 e4. doi: 10.1016/j.cub.2024.12.012.

24. Bettencourt KC, Xu Y. Decoding the content of visual short-term memory under distraction in occipital and parietal areas. Nat Neurosci. 2016;19(1):150–7. doi: 10.1038/nn.4174.

25. Rademaker RL, Chunharas C, Serences JT. Coexisting representations of sensory and mnemonic information in human visual cortex. Nat Neurosci. 2019;22(8):1336–44. doi: 10.1038/s41593-019-0428-x.

26. Bosch SE, Jehee JF, Fernandez G, Doeller CF. Reinstatement of associative memories in early visual cortex is signaled by the hippocampus. J Neurosci. 2014;34(22):7493–500. doi: 10.1523/JNEUROSCI.0805-14.2014.

27. Hu Y, Yu Q. Spatiotemporal dynamics of self-generated imagery reveal a reverse cortical hierarchy from cue-induced imagery. Cell Rep. 2023;42(10):113242. doi: 10.1016/j.celrep.2023.113242.

28. Li S, Zeng X, Shao Z, Yu Q. Neural Representations in Visual and Parietal Cortex Differentiate between Imagined, Perceived, and Illusory Experiences. J Neurosci. 2023;43(38):6508–24. doi: 10.1523/JNEUROSCI.0592-23.2023.

29. Albers AM, Kok P, Toni I, Dijkerman HC, de Lange FP. Shared representations for working memory and mental imagery in early visual cortex. Curr Biol. 2013;23(15):1427–31. doi: 10.1016/j.cub.2013.05.065.

30. Vo VA, Sutterer DW, Foster JJ, Sprague TC, Awh E, Serences JT. Shared Representational Formats for Information Maintained in Working Memory and Information Retrieved from Long-Term Memory. Cereb Cortex. 2022;32(5):1077–92. doi: 10.1093/cercor/bhab267.

31. Favila SE, Kuhl BA, Winawer J. Perception and memory have distinct spatial tuning properties in human visual cortex. Nat Commun. 2022;13(1):5864. doi: 10.1038/s41467-022-33161-8.

32. Lawrence SJD, van Mourik T, Kok P, Koopmans PJ, Norris DG, de Lange FP. Laminar Organization of Working Memory Signals in Human Visual Cortex. Curr Biol. 2018;28(21):3435–40 e4. doi: 10.1016/j.cub.2018.08.043.

33. Iamshchinina P, Kaiser D, Yakupov R, Haenelt D, Sciarra A, Mattern H, et al. Perceived and mentally rotated contents are differentially represented in cortical depth of V1. Commun Biol. 2021;4(1):1069. doi: 10.1038/s42003-021-02582-4.

34. Mechelli A, Price CJ, Friston KJ, Ishai A. Where bottom-up meets top-down: neuronal interactions during perception and imagery. Cereb Cortex. 2004;14(11):1256–65. doi: 10.1093/cercor/bhh087.

35. Pearson J. The human imagination: the cognitive neuroscience of visual mental imagery. Nat Rev Neurosci. 2019;20(10):624–34. doi: 10.1038/s41583-019-0202-9.

36. Hochstein S, Ahissar M. View from the top: hierarchies and reverse hierarchies in the visual system. Neuron. 2002;36(5):791–804. doi: 10.1016/s0896-6273(02)01091-7.

37. Dijkstra N, Zeidman P, Ondobaka S, van Gerven MAJ, Friston K. Distinct Top-down and Bottom-up Brain Connectivity During Visual Perception and Imagery. Sci Rep. 2017;7(1):5677. doi: 10.1038/s41598-017-05888-8.

38. Shi D, Yu Q. Distinct neural signatures underlying information maintenance and manipulation in working memory. Cereb Cortex. 2024;34(3). doi: 10.1093/cercor/bhae063.

39. Miller EK, Cohen JD. An integrative theory of prefrontal cortex function. Annu Rev Neurosci. 2001;24:167–202. doi: 10.1146/annurev.neuro.24.1.167.

40. Breedlove JL, St-Yves G, Olman CA, Naselaris T. Generative Feedback Explains Distinct Brain Activity Codes for Seen and Mental Images. Current Biology. 2020;30(12):2211–24.e6. doi: 10.1016/j.cub.2020.04.014.

41. Xu Y. Parietal-driven visual working memory representation in occipito-temporal cortex. Curr Biol. 2023;33(20):4516–23 e5. doi: 10.1016/j.cub.2023.08.080.

42. Liebe S, Hoerzer GM, Logothetis NK, Rainer G. Theta coupling between V4 and prefrontal cortex predicts visual short-term memory performance. Nat Neurosci. 2012;15(3):456–62, S1-2. doi: 10.1038/nn.3038.

43. Zhang M, Yu Q. The representation of abstract goals in working memory is supported by task-congruent neural geometry. PLoS Biol. 2024;22(12):e3002461. doi: 10.1371/journal.pbio.3002461.

44. Pearson J, Westbrook F. Phantom perception: voluntary and involuntary nonretinal vision. Trends Cogn Sci. 2015;19(5):278–84. doi: 10.1016/j.tics.2015.03.004.

45. Dawes AJ, Keogh R, Andrillon T, Pearson J. A cognitive profile of multi-sensory imagery, memory and dreaming in aphantasia. Sci Rep. 2020;10(1):10022. doi: 10.1038/s41598-020-65705-7.

46. Dawes AJ, Keogh R, Pearson J. Multisensory subtypes of aphantasia: Mental imagery as supramodal perception in reverse. Neurosci Res. 2024;201:50–9. doi: 10.1016/j.neures.2023.11.009.

47. Liu S, Yu Q, Tse PU, Cavanagh P. Neural Correlates of the Conscious Perception of Visual Location Lie Outside Visual Cortex. Curr Biol. 2019;29(23):4036–44 e4. doi: 10.1016/j.cub.2019.10.033.

48. Lisi M, Cavanagh P. Dissociation between the Perceptual and Saccadic Localization of Moving Objects. Curr Biol. 2015;25(19):2535–40. doi: 10.1016/j.cub.2015.08.021.

49. Yu Q, Postle BR. The Neural Codes Underlying Internally Generated Representations in Visual Working Memory. J Cogn Neurosci. 2021:1–16. doi: 10.1162/jocn_a_01702.

50. Spagna A, Hajhajate D, Liu J, Bartolomeo P. Visual mental imagery engages the left fusiform gyrus, but not the early visual cortex: A meta-analysis of neuroimaging evidence. Neurosci Biobehav Rev. 2021;122:201–17. doi: 10.1016/j.neubiorev.2020.12.029.

51. Liu J, Zhan M, Hajhajate D, Spagna A, Dehaene S, Cohen L, et al. Visual mental imagery in typical imagers and in aphantasia: A millimeter-scale 7-T fMRI study. Cortex. 2025;185:113–32. doi: 10.1016/j.cortex.2025.01.013.

52. Liu J, Bartolomeo P. Aphantasia as a functional disconnection. Trends Cogn Sci. 2025;29(11):963–4. doi: 10.1016/j.tics.2025.05.012.

53. Kwak Y, Curtis CE. Unveiling the abstract format of mnemonic representations. Neuron. 2022;110(11):1822–8 e5. doi: 10.1016/j.neuron.2022.03.016.

54. Spaak E, Watanabe K, Funahashi S, Stokes MG. Stable and Dynamic Coding for Working Memory in Primate Prefrontal Cortex. J Neurosci. 2017;37(27):6503–16. doi: 10.1523/JNEUROSCI.3364-16.2017.

55. Yu Q, Shim WM. Temporal-Order-Based Attentional Priority Modulates Mnemonic Representations in Parietal and Frontal Cortices. Cereb Cortex. 2019;29(7):3182–92. doi: 10.1093/cercor/bhy184.

56. Kriegeskorte N, Mur M, Bandettini P. Representational similarity analysis - connecting the branches of systems neuroscience. Front Syst Neurosci. 2008;2:4. doi: 10.3389/neuro.06.004.2008.

57. McLaren DG, Ries ML, Xu G, Johnson SC. A generalized form of context-dependent psychophysiological interactions (gPPI): a comparison to standard approaches. Neuroimage. 2012;61(4):1277–86. doi: 10.1016/j.neuroimage.2012.03.068.

58. Frey M, Nau M, Doeller CF. Magnetic resonance-based eye tracking using deep neural networks. Nat Neurosci. 2021;24(12):1772–9. doi: 10.1038/s41593-021-00947-w.

59. Yu Q, Panichello MF, Cai Y, Postle BR, Buschman TJ. Delay-period activity in frontal, parietal, and occipital cortex tracks noise and biases in visual working memory. PLoS Biol. 2020;18(9):e3000854. doi: 10.1371/journal.pbio.3000854.

60. Dake M, Curtis CE. Perturbing human V1 degrades the fidelity of visual working memory. Nat Commun. 2025;16(1):2675. doi: 10.1038/s41467-025-57882-8.

61. Leavitt ML, Mendoza-Halliday D, Martinez-Trujillo JC. Sustained Activity Encoding Working Memories: Not Fully Distributed. Trends Neurosci. 2017;40(6):328–46. doi: 10.1016/j.tins.2017.04.004.

62. Huang J, Wang T, Dai W, Li Y, Yang Y, Zhang Y, et al. Neuronal representation of visual working memory content in the primate primary visual cortex. Sci Adv. 2024;10(24):eadk3953. doi: 10.1126/sciadv.adk3953.

63. Li HH, Ma WJ, Curtis CE. Dynamics of working memory drift and information flow across the cortical hierarchy. Proc Natl Acad Sci U S A. 2026;123(4):e2518110123. doi: 10.1073/pnas.2518110123.

64. Li HH, Curtis CE. Neural population dynamics of human working memory. Curr Biol. 2023;33(17):3775–84 e4. doi: 10.1016/j.cub.2023.07.067.

65. Monzel M, Leelaarporn P, Lutz T, Schultz J, Brunheim S, Reuter M, et al. Hippocampal-occipital connectivity reflects autobiographical memory deficits in aphantasia. Elife. 2024;13. doi: 10.7554/eLife.94916.

66. Montabes de la Cruz BM, Abbatecola C, Luciani RS, Paton AT, Bergmann J, Vetter P, et al. Decoding sound content in the early visual cortex of aphantasic participants. Curr Biol. 2024;34(21):5083–9 e3. doi: 10.1016/j.cub.2024.09.008.

67. Milton F, Fulford J, Dance C, Gaddum J, Heuerman-Williamson B, Jones K, et al. Behavioral and Neural Signatures of Visual Imagery Vividness Extremes: Aphantasia versus Hyperphantasia. Cereb Cortex Commun. 2021;2(2):tgab035. doi: 10.1093/texcom/tgab035.

68. Ragni F, Tucciarelli R, Andersson P, Lingnau A. Decoding stimulus identity in occipital, parietal and inferotemporal cortices during visual mental imagery. Cortex. 2020;127:371–87. doi: 10.1016/j.cortex.2020.02.020.

69. Krempel R, Monzel M. Aphantasia and involuntary imagery. Conscious Cogn. 2024;120:103679. doi: 10.1016/j.concog.2024.103679.

70. Bergmann J, Ortiz-Tudela J. Feedback signals in visual cortex during episodic and schematic memory retrieval and their potential implications for aphantasia. Neurosci Biobehav Rev. 2023;152:105335. doi: 10.1016/j.neubiorev.2023.105335.

71. Marks DF. Visual imagery differences in the recall of pictures. Br J Psychol. 1973;64(1):17–24. doi: 10.1111/j.2044-8295.1973.tb01322.x.

72. Reisberg D, Pearson DG, Kosslyn SM. Intuitions and introspections about imagery: the role of imagery experience in shaping an investigator’s theoretical views. Applied Cognitive Psychology. 2003;17(2):147–60. doi: 10.1002/acp.858.

73. Andrade J, May J, Deeprose C, Baugh SJ, Ganis G. Assessing vividness of mental imagery: The Plymouth Sensory Imagery Questionnaire. Br J Psychol. 2014;105(4):547–63. doi: 10.1111/bjop.12050.

74. Nystrom M, Holmqvist K. An adaptive algorithm for fixation, saccade, and glissade detection in eyetracking data. Behav Res Methods. 2010;42(1):188–204. doi: 10.3758/BRM.42.1.188.

75. Cox RW. AFNI: Software for Analysis and Visualization of Functional Magnetic Resonance Neuroimages. Computers and Biomedical Research. 1996;29(3):162–73. doi: 10.1006/cbmr.1996.0014.

76. Wang L, Mruczek RE, Arcaro MJ, Kastner S. Probabilistic Maps of Visual Topography in Human Cortex. Cereb Cortex. 2015;25(10):3911–31. doi: 10.1093/cercor/bhu277.

77. Desikan RS, Segonne F, Fischl B, Quinn BT, Dickerson BC, Blacker D, et al. An automated labeling system for subdividing the human cerebral cortex on MRI scans into gyral based regions of interest. Neuroimage. 2006;31(3):968–80. doi: 10.1016/j.neuroimage.2006.01.021.

